# Automated assembly of high-quality diploid human reference genomes

**DOI:** 10.1101/2022.03.06.483034

**Authors:** Erich D. Jarvis, Giulio Formenti, Arang Rhie, Andrea Guarracino, Chentao Yang, Jonathan Wood, Alan Tracey, Francoise Thibaud-Nissen, Mitchell R. Vollger, David Porubsky, Haoyu Cheng, Mobin Asri, Glennis A. Logsdon, Paolo Carnevali, Mark J.P. Chaisson, Chen-Shan Chin, Sarah Cody, Joanna Collins, Peter Ebert, Merly Escalona, Olivier Fedrigo, Robert S. Fulton, Lucinda L. Fulton, Shilpa Garg, Jay Ghurye, Ana Granat, Edward Green, Ira Hall, William Harvey, Patrick Hasenfeld, Alex Hastie, Marina Haukness, Erich B. Jaeger, Miten Jain, Melanie Kirsche, Mikhail Kolmogorov, Jan O. Korbel, Sergey Koren, Jonas Korlach, Joyce Lee, Daofeng Li, Tina Lindsay, Julian Lucas, Feng Luo, Tobias Marschall, Jennifer McDaniel, Fan Nie, Hugh E. Olsen, Nathan D. Olson, Trevor Pesout, Daniela Puiu, Allison Regier, Jue Ruan, Steven L. Salzberg, Ashley D. Sanders, Michael C. Schatz, Anthony Schmitt, Valerie A. Schneider, Siddarth Selvaraj, Kishwar Shafin, Alaina Shumate, Catherine Stober, James Torrance, Justin Wagner, Jianxin Wang, Aaron Wenger, Chuanle Xiao, Aleksey V. Zimin, Guojie Zhang, Ting Wang, Heng Li, Erik Garrison, David Haussler, Justin M. Zook, Evan E. Eichler, Adam M. Phillippy, Benedict Paten, Kerstin Howe, Karen H. Miga, Human Pangenome Reference Consortium

## Abstract

The current human reference genome, GRCh38, represents over 20 years of effort to generate a high-quality assembly, which has greatly benefited society^1, 2^. However, it still has many gaps and errors, and does not represent a biological human genome since it is a blend of multiple individuals^3, 4^. Recently, a high-quality telomere-to-telomere reference genome, CHM13, was generated with the latest long-read technologies, but it was derived from a hydatidiform mole cell line with a duplicate genome, and is thus nearly homozygous^5^. To address these limitations, the Human Pangenome Reference Consortium (HPRC) recently formed with the goal of creating a collection of high-quality, cost-effective, diploid genome assemblies for a pangenome reference that represents human genetic diversity^6^. Here, in our first scientific report, we determined which combination of current genome sequencing and automated assembly approaches yields the most complete, accurate, and cost-effective diploid genome assemblies with minimal manual curation. Approaches that used highly accurate long reads and parent-child data to sort haplotypes during assembly outperformed those that did not. Developing a combination of all the top performing methods, we generated our first high-quality diploid reference assembly, containing only ∼4 gaps (range 0-12) per chromosome, most within ± 1% of CHM13’s length. Nearly 1/4th of protein coding genes have synonymous amino acid changes between haplotypes, and centromeric regions showed the highest density of variation. Our findings serve as a foundation for assembling near-complete diploid human genomes at the scale required for constructing a human pangenome reference that captures all genetic variation from single nucleotides to large structural rearrangements.

## Main

The initial draft of the current human reference genome was the main outcome of over a decade of effort by the Human Genome Project (HGP), with total cost exceeding $2.7 billion (over $5 billion at today’s value)^1, 2, 7^. Its current build, GRCh38, reflects two decades of additional effort by the Genome Reference Consortium and others to correct the initial draft. It was created from physical maps of thousands of individually sequenced 40-2,000 kb BAC, YAC, and fosmid clones, supplemented with whole genome sequence data^1, 2^. It is also a combination of DNA sequences from 20 anonymous volunteers, with one individual representing ∼70% of the sequence^2^. Over the years, GRCh38 was advanced from having over 150,000 gaps to currently 995 in the primary assembly^2, 3^. Therefore, despite being one of the most complete human reference genomes available, GRCh38 represents an incomplete composite, and does not adequately capture the full spectrum of humankind genomic variation^8^.

In the years following the HGP, several technological limitations prevented the generation of new human reference genomes of similar or higher quality at scale. Sequence duplications much larger than the sequence read lengths are particularly challenging to assemble. The sequencing enzymes used often have difficulty reading through regions with complex structures, such as GC-rich regions often found in promoters that regulate gene expression^9, 10^. It is also now clear that mixing diverse haplotypes in a single assembly, even from the same individual, can introduce many errors with standard assembly tools^8, 10, 11^. These errors include: switch errors where variants from each haplotype are assembled into the same pseudo-haplotype; false duplications and associated gaps where more divergent haplotype homologs are assembled as separate false paralogs; and consensus errors due to collapses between haplotypes. While resequencing efforts using less expensive short reads have contributed to revealing more single-nucleotide variation, structural variation is not fully captured^10, 13^.

Major improvements have since been made in sequence read lengths^4, 14^, long read base accuracy^15^, contig algorithms, scaffolding contigs into chromosomes^9, 16–18^, haplotype resolution^19–21^, and technologies with reduced sequencing cost. These advances include those made by the Vertebrate Genomes Project (VGP)^9^, the Human Genome Structural Variation Consortium (HGSVC)^13^, and the Telomere-to-Telomere (T2T) consortium, the latter of which produced the first complete human reference genome, of CHM13^5^. The CHM13 cell line originated from a hydatidiform mole with nearly two identical paternal haploid complements inherited from the maternal line, eliminating the need to separate haplotypes and the associated diploid assembly errors. Completing the T2T CHM13 assembly also required a significant amount of manual curation by dozens of persons over many months, with different groups focused on each chromosome. Thus, despite improvements, additional developments are needed to assemble diploid genomes at high-quality and at scale, which are critical for clinically relevant samples and understanding human genetic variation.

To help overcome these limitations, in 2019 the National Human Genome Research Institute (NHGRI) invested in an international Human Pangenome Reference Consortium (HPRC), with an aspired goal of producing a high-quality pangenome reference representing over 99% of human genetic diversity for minor alleles of 1% or higher frequency in the human population^6^. We estimate that one could start to approach this goal with complete *de novo* assemblies of ∼450 individuals (e.g. 900 haplotypes) from the world population (**Supplementary Note 1**). Starting in 2020, we tested the current best practices in sequencing technologies and automated assembly algorithms on one human sample, HG002, an openly- consented Ashkenazi individual from the Personal Genome Project^22^. We included parental samples (HG003-father, HG004-mother) for trio-based assemblies, where parental sequence data were used to sort haplotypes in the offspring sequence data^9, 20^. Extensive evaluation of the resulting assemblies alongside GRCh38 and CMH13 led to new approaches that yielded the best of over 60 metrics, and new biological discoveries. Evaluation also identified areas of needed improvement to achieve automated complete and error-free diploid genome assemblies.

### Data types and algorithms

We chose HG002 because of available prior extensive public data^23^ and benchmarks^24^ generated by the Genome in a Bottle (GIAB) consortium. As a male sample, it also enables the assembly and evaluation of both X and Y chromosomes. We obtained or generated additional state-of-the-art sequence data types, including Pacbio HiFi long reads and Oxford Nanopore (ONT) long reads (>10 kb) for generating contigs, and long-range link information (e.g. 10X linked reads, Hi-C linked reads, optical maps, and Strand-seq) for scaffolding the contigs (**Supplementary Table 1**). These choices were made based on lessons learned for producing high-quality assemblies from other consortia (e.g. VGP^9^, T2T^5^, HGSVC^13^) or individual labs^25–27^. In particular, long-read based assemblies are more contiguous and more structurally accurate than short-read based assemblies, long-range linking information can place contigs into chromosome-level scaffolds, and haplotype phasing and high base accuracy helps prevent false duplications and other common assembly errors.

We made an open call to the international genome community to produce the most complete and highest-quality genome assembly possible of HG002 with the data provided, in an automated manner without manual curation (https://humanpangenome.org/hg002/). We received 23 assembly combinations, from 14 groups, including HPRC members, that used different data types and algorithms for contig assembly, scaffolding, and/or haplotype phasing when attempted (**Table 1**); we named them asm1 to asm23, with suffixes a/b for haplotypes. Among these 23, 12 assembly algorithms were used: Canu and HiCanu^15^, CrossStitch, DipAsm^27^, FALCON Unzip^19^, Flye^28^, hifiasm^29^, MaSuRCA^30^, NECAT^31^, Peregrine^32^, Shasta^26^, and wtdbg^33^ (**Table 1**). We classified the assemblies into four categories: 1) diploid scaffolded assemblies, where there was an attempt to assemble comparable contigs and scaffolds of both haplotypes or two pseudohaplotypes (mixed paternal and maternal derived sequences); 2) diploid contig-only assemblies, where there was an attempt to assemble comparable contigs only of both haplotypes/pseudohaplotypes or a more complete assembly representing one pseudohaplotype; 3) haploid scaffolded assemblies, where contigs and scaffolds were all merged into one pseudohaplotype; and 4) haploid contig-only assemblies, where only contigs were generated and all merged into one pseudohaplotype (**Table 1, Supplementary Table 2a,b**). CrossStitch is reference-based (to GRCh38 in this study) by design, and MaSuRCA used GRCh38 to order and orient assembled HG002 contigs into chromosome-level scaffolds, followed by filling in gaps with GRCh38 sequence. Since these assemblies (asm1, asm15, and asm17) are not “pure” *de novo*, they might contain some biases towards the reference, certainly for gap filling. They are included for metric comparisons and to establish a baseline for capturing variation guided by a reference genome, but are considered cautiously for *de novo* methods going forward. Following the VGP model^9^, we assessed over 60 metrics under 14 categories, including continuity, structural accuracy, base accuracy, haplotype phasing, and completeness (**Supplementary Table 2**).

**Table 1.**
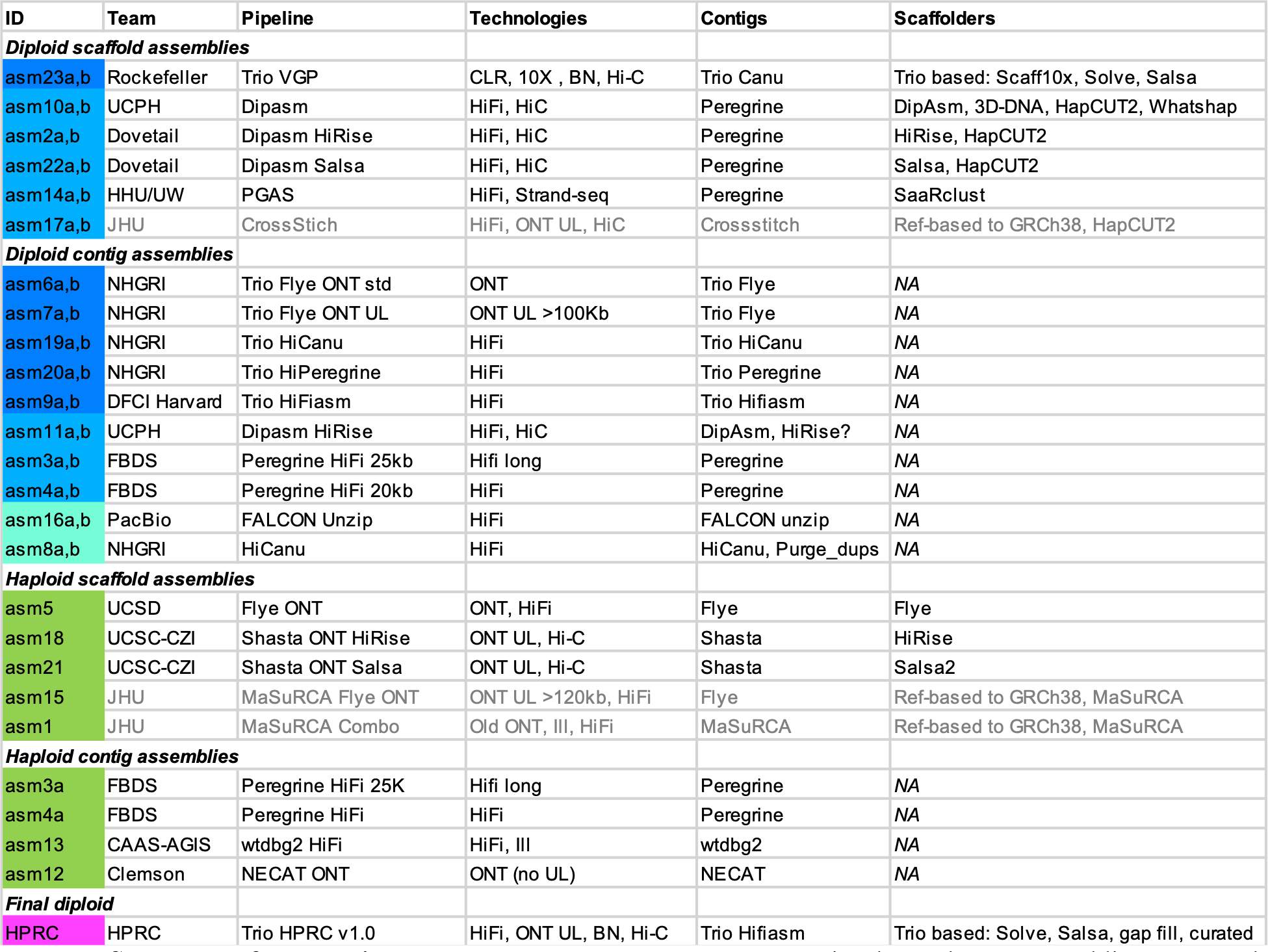
Summary of sequencing and assembly approaches tested. Listed are the 23 assemblies generated, categorized into four broad types based on whether there were diploid (a,b) or merged haploid, and scaffolded or contigs only. They are color coded according to type of haplotype phasing: blue, trio-based; light blue, non-trio and more complete for both haplotypes; turquoise, non-trio and more complete for only one haplotype; green, a merging of haplotypes. The final HPRC reference v1.0 is colored magenta to distinguish it from the test assemblies, but is trio-based. Grey-text entries, referenced-based assemblies. More details on the sequencing technologies are in Supplementary Table 1, and the contig and scaffold assemblers in the methods and Supplementary Table 2a,b.

| ID | Team | Pipeline | Technologies | Contigs | Scaffolders |
| --- | --- | --- | --- | --- | --- |
| **Diploid scaffold assemblies** |  |  |  |  |  |
| asm23a,b | Rockefeller | Trio VGP | CLR, 10X , BN, Hi-C | Trio Canu | Trio based: Scaff10x, Solve, Salsa |
| asm10a,b | UCPH | Dipasm | HiFi, HiC | Peregrine | DipAsm, 3D-DNA, HapCUT2, Whatshap |
| asm2a,b | Dovetail | Dipasm HiRise | HiFi, HiC | Peregrine | HiRise, HapCUT2 |
| asm22a,b | Dovetail | Dipasm Salsa | HiFi, HiC | Peregrine | Salsa, HapCUT2 |
| asm14a,b | HHU/UW | PGAS | HiFi, Strand-seq | Peregrine | SaaRclust |
| asm17a,b | JHU | CrossStitch | HiFi, ONT UL, HiC | Crossstitch | Ref-based to GRCh38, HapCUT2 |
| **Diploid contig assemblies** |  |  |  |  |  |
| asm6a,b | NHGRI | Trio Flye ONT std | ONT | Trio Flye | NA |
| asm7a,b | NHGRI | Trio Flye ONT UL | ONT UL >100Kb | Trio Flye | NA |
| asm19a,b | NHGRI | Trio HiCanu | HiFi | Trio HiCanu | NA |
| asm20a,b | NHGRI | Trio HiPeregrine | HiFi | Trio Peregrine | NA |
| asm9a,b | DFCI Harvard | Trio HiFiasm | HiFi | Trio Hifiasm | NA |
| asm11a,b | UCPH | Dipasm HiRise | HiFi, HiC | DipAsm, HiRise? | NA |
| asm3a,b | FBDS | Peregrine HiFi 25kb | Hifi long | Peregrine | NA |
| asm4a,b | FBDS | Peregrine HiFi 20kb | HiFi | Peregrine | NA |
| asm16a,b | PacBio | FALCON Unzip | HiFi | FALCON unzip | NA |
| asm8a,b | NHGRI | HiCanu | HiFi | HiCanu, Purge_dups | NA |
| **Haploid scaffold assemblies** |  |  |  |  |  |
| asm5 | UCSD | Flye ONT | ONT, HiFi | Flye | Flye |
| asm18 | UCSC-CZI | Shasta ONT HiRise | ONT UL, Hi-C | Shasta | HiRise |
| asm21 | UCSC-CZI | Shasta ONT Salsa | ONT UL, Hi-C | Shasta | Salsa2 |
| asm15 | JHU | MaSuRCA Flye ONT | ONT UL >120kb, HiFi | Flye | Ref-based to GRCh38, MaSuRCA |
| asm1 | JHU | MaSuRCA Combo | Old ONT, III, HiFi | MaSuRCA | Ref-based to GRCh38, MaSuRCA |
| **Haploid contig assemblies** |  |  |  |  |  |
| asm3a | FBDS | Peregrine HiFi 25K | Hifi long | Peregrine | NA |
| asm4a | FBDS | Peregrine HiFi | HiFi | Peregrine | NA |
| asm13 | CAAS-AGIS | wtdbg2 HiFi | HiFi, III | wtdbg2 | NA |
| asm12 | Clemson | NECAT ONT | ONT (no UL) | NECAT | NA |
| **Final diploid** |  |  |  |  |  |
| HPRC | HPRC | Trio HPRC v1.0 | HiFi, ONT UL, BN, Hi-C | Trio Hifiasm | Trio based: Solve, Salsa, gap fill, curated |

### Non-human contamination and organelle genomes

First, we screened for non-human DNA and found that all *de novo* assemblies had between 1 to 25 contigs or scaffolds with library adaptor sequence contamination, which were not successfully removed during read preprocessing (**Extended Data Fig. 1a**, **Supplementary Table 2c**). The presence of adaptor sequences on reads with human sequences introduced gaps between the human-based contigs; reads with adaptor alone were concatenated to make adaptor only contigs (**Supplementary Note 2**). We also found instances of assembled bacterial (*Escherichia coli*) and yeast (*Saccharomyces cerevisiae*) genomes, either as stand- alone contigs or scaffolds (3 assemblies), chimeric with human genomic DNA (4 assemblies), or both (4 assemblies; **Extended Data Fig. 1b**, **Supplementary Table 2c**). There were typically 0 to 6 copies of these microbial genomes per assembly, except in the wtdgb2 assembly with 35 *E. coli* and 46 *S. cerevisiae* contigs. There were also from 1 to ∼ 40 assembled human mitochondrial (MT) contigs in ∼74% (17/23) of the assemblies (**Extended Data Fig. 1c, Supplementary Table 2c**). In the trio-based assemblies, they were all associated with the maternal haplotype, indicating that the MT reads were correctly sorted during haplotype phasing before generating contigs (in the VGP Trio assembly the MT was purposely included in both haplotypes to avoid NUMT overpolishing^9^). Most mitochondrial contigs were full length genomes, further demonstrating^34^ that with long reads most new assembly algorithms can assemble a mitochondrial genome in one contig. Part of the reason for the differential presence of MT genomes in the assemblies is presumably due to differential read length thresholds used for initial contig assembly; the higher the size threshold, the less likely MT reads will be included^34^.

**Fig. 1.**
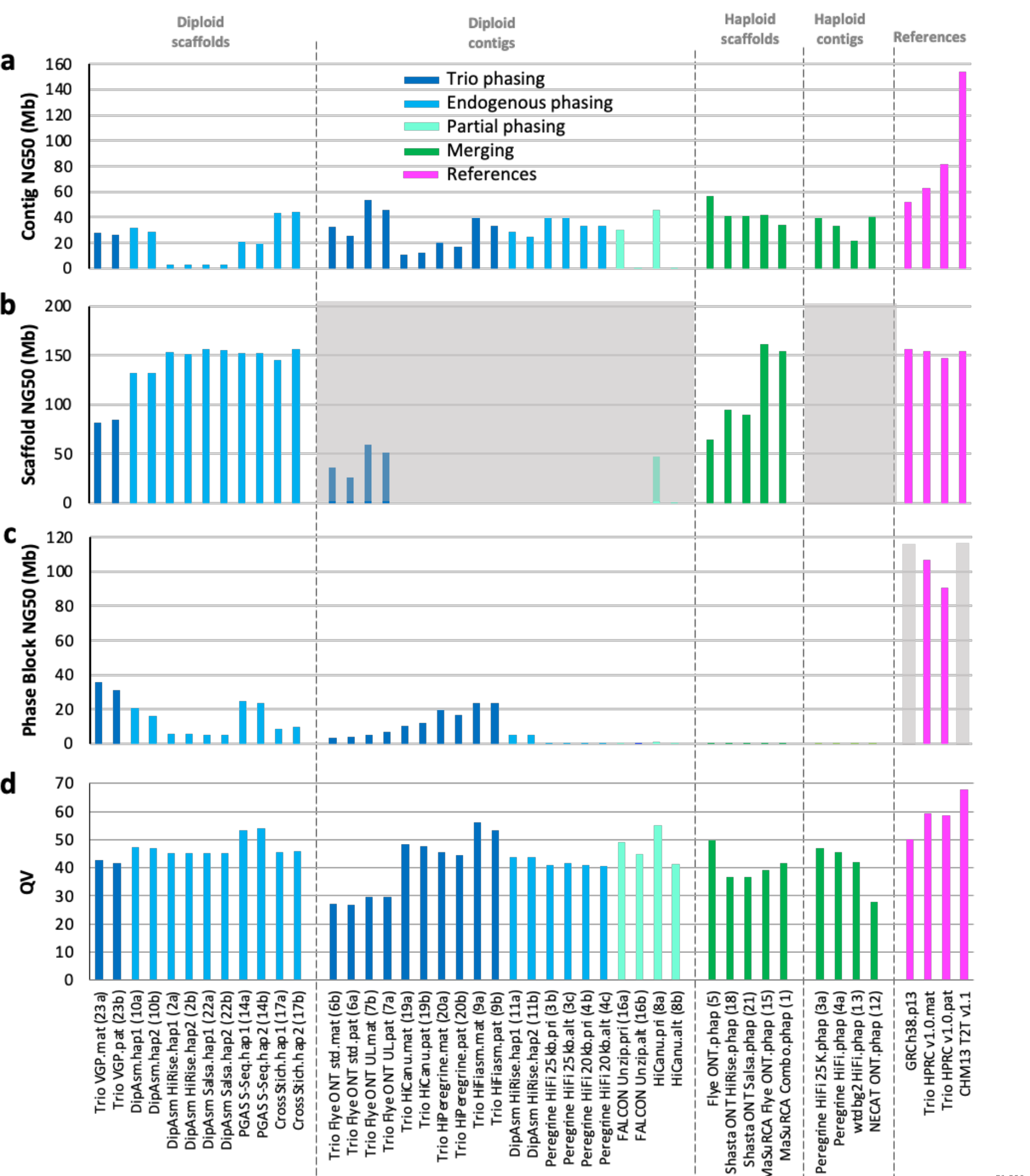
Assembly continuity, phasing, and base call accuracy metrics. **a,** Contig NG50 values**. b,** Scaffold NG50 values**. c**, Haplotype phase block NG50 values. **d**, QV base call accuracy; as an example, QV60 is about 1 error per Mb. Color code designates type of haplotype phasing performed. Grey shaded regions in (b) are not applicable for scaffold metrics, as these are contig-only assemblies; however, several contig only assemblies had gaps, because the Flye assembler inserts gaps in contigs where there is uncertainty of a repeat sequence; and the purge_dups function applied to the HiCanu contigs removes false duplications within contigs and creates a gap in the removed location. Grey shading in (c) indicates not applicable for phase blocks, because the GRCh38 has many haplotypes and CHM13 is from a haploid cell line. () along the x-axis are the assembly (asm) numbers.

### Highly contiguous phased assemblies

Our assembly targets were an expected maternal genome size of ∼3.06 Gb (22 autosomes + X) and paternal size of ∼2.96 Gb (22 autosomes + Y), given the expected X (155.3 Mb) and Y (∼60 Mb) difference of about ∼96 Mb^35^. Almost all assemblies, including the diploid ones, were close to the expected sizes of a human genome (∼3.0 Gb; range 2.8-3.1 Gb; **Extended Data Fig. 2a-c, Supplementary Table 2d-f**). Only the diploid pair asm19a and asm19b were bigger, by ∼3%. In the trio-based assemblies, the maternal (mat) haplotype were all longer than the paternal (pat), consistent with sex chromosome differences. In the non-trio diploid-based assemblies, each haplotype was more similar in length and tended to be skewed toward the expected size of the maternal haplotype, consistent with our finding of either both X and part of Y in both haplotypes, or missing the Y assembly altogether (**Supplementary Table 2d,** assessed for the diploid scaffolded assemblies). The assemblies that came closest to the theoretical expected total size (98-100%) for both maternal and paternal haplotypes were the Trio VGP scaffolded (asm23a,b) and the Trio hifiasm (asm9a,b) assemblies (**Extended Data Fig. 2a**). Although close to the expected sizes, the scaffolded assemblies still had quite a range of missing sequence (Ns), from ∼40 kb to 50 Mb, either in the gaps between contigs or trailing Ns at scaffold ends (**Extended Data Fig. 2c; Supplementary Table 2f**). In comparison, GRCh38 has ∼151 Mb of gaps. We note that with the exception of Bionano optical maps, most scaffolding tools place arbitrary gap sizes. Most assemblies also had between 0.3-2.3% false duplications (according to kmer counts; **Extended Data Fig. 2d, Supplementary Table 2g**), which could also explain why some were bigger than expected^9^. GRCh38 also still contains false duplications^5, 36^, although they are difficult to estimate precisely due to the mixture of haplotypes.

**Figure 2.**
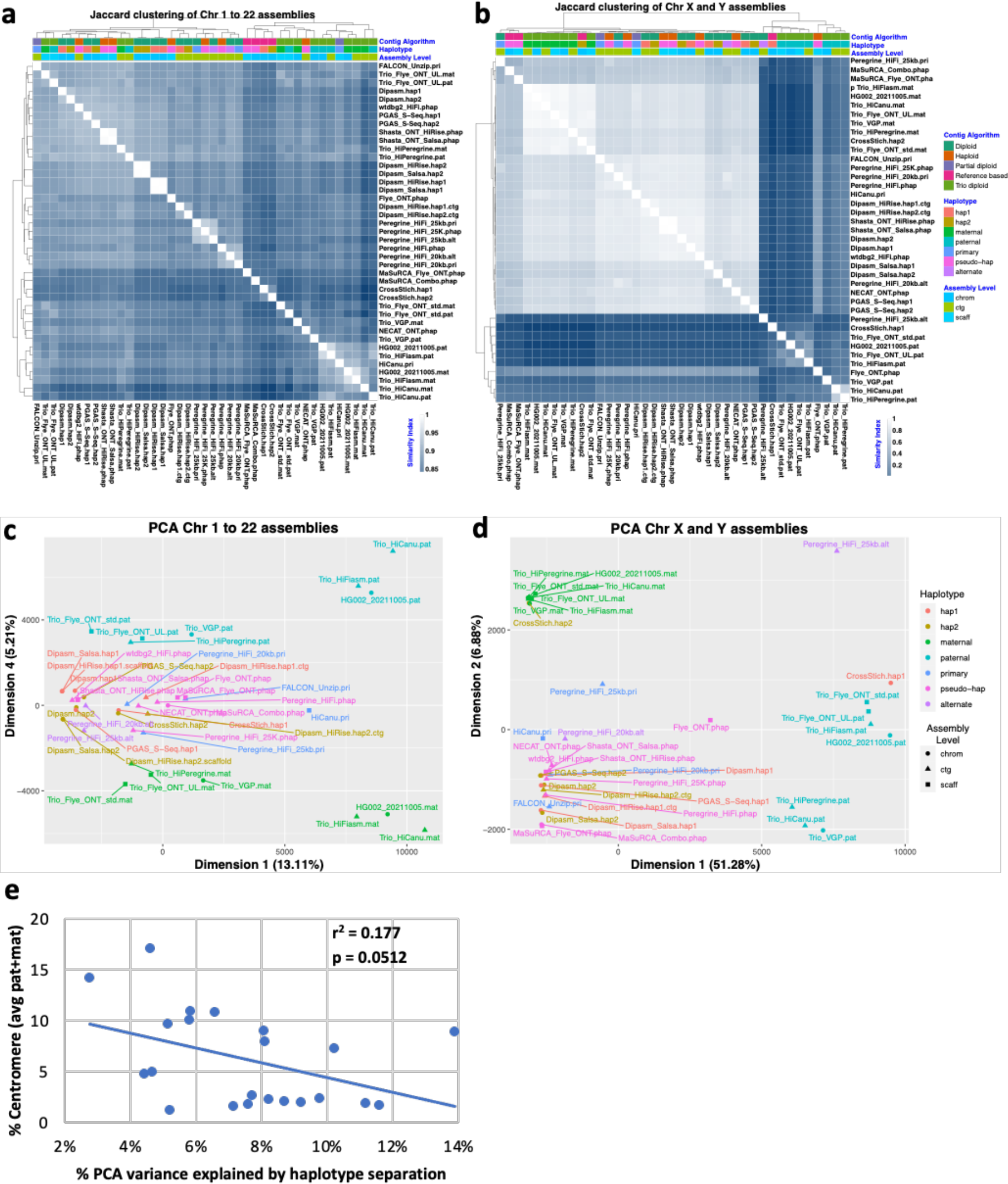
Multidimensional relationship among assemblies. **a** and **b**, Clustering of pairwise Jaccard similiarities between pairs of assemblies, for the autosomes 1-22 and sex chromosomes X and Y, respectively. Heatmap, the lighter the blue (closer to 1) the more similarity between pairs of assemblies. Assemblies are annotated with four different color-coded classifications. **c** and **d**, Principle component analysis on the multidimensional Euclidean distances amongst assemblies, for the autosomes 1-22 and sex chromosomes X and Y, respectively. PCA dimensions shown are those where the paternal and maternal haplotypes separated the strongest. **e**, Correlation between % centromere:chromosome size and % PCA variance where trio-based autosome assemblies separated by haplotype.

In terms of continuity, our goal was to minimize the number of gaps for a theoretical maximum gapless contig NG50 that equals a chromosome NG50 of ∼155 Mb for human (where half of the assembled contigs are this size and bigger)^5^. Most assemblies had contig NG50 sizes in the 20-50 Mb range, including for both haplotypes for some of the diploid assemblies (**Fig. 1a; Supplementary Table 2e**). These values are between ∼13-32% of the theoretical maximum of 155 Mb for the maternal haplotype, indicating partial chromosomal length contigs. Exceptions well below NG50 of 20 Mb were: the alternative (alt) haplotypes of FALCON Unzip or HiCanu approaches that generate a partial diploid assembly by design (asm16b and asm8b, respectively), with the primary pseudohaplotype being more contiguous (asm16a and asm8a); and both haplotypes of the Dovetail implementation of the DipAsm assembler (asm2 and asm22), where Hi-C data was used to phase the haplotypes. In contrast, the original implementation of DipAsm created two assemblies with contig NG50s greater than 20 Mb (asm10a,b). Not surprisingly, the assembly (asm7a,b) that used the ONT ultralong reads (ONT UL > 100 kb) had the highest contig NG50s (48.6 Mb maternal; 39.8 Mb paternal). The trio-based ONT and hifiasm (asm9a,b) HiFi assemblies had the fewest contigs (∼600-900) of all diploid assemblies (**Extended Data Fig. 3a**). All scaffolded assemblies had scaffold NG50 values ranging from 80-155 Mb (**Fig. 1b**), 52-100% of the theoretical maximum of 155 Mb. All non-trio diploid scaffolded assemblies had 23-30 scaffolds, at or close to the expected 23 chromosomes per haplotype (**Supplementary Table 2f**). However, this particular metric comparison is made less informative since DipAsm inherently filters out scaffolds less than 10 kb, PGAS excludes contigs less than 500 kb as the Strand-seq signal is too sparse to scaffold small contigs, and CrossStich only includes contigs/scaffolds that align to the GRCh38 reference. Such exclusions have the potential to result in higher numbers of gaps with missing sequence. This idea is supported by the Trio VGP scaffolded assembly (asm23a,b) that did not exclude scaffolds based on size or alignment to a reference, where, not surprisingly, it had a much higher number of scaffolds (over 2,000 each) but fewer gaps among those scaffolds (673 maternal and 917 paternal) relative to DipAsm and PGAS assemblies (900-4,000 within scaffold gaps; **Extended Data Fig. 3b,c**). The size of the largest scaffold (max) for most assemblies approached the size of chromosome 1 (248 Mb; range 149-242 Mb; **Supplementary Table 2f**). Taken together, these findings demonstrate an important shift in recent assembly tools to generate two separate assemblies per individual, representing the two haplotypes or pseudohaplotypes, with chromosome-level scaffolds, albeit with gaps.

**Fig. 3.**
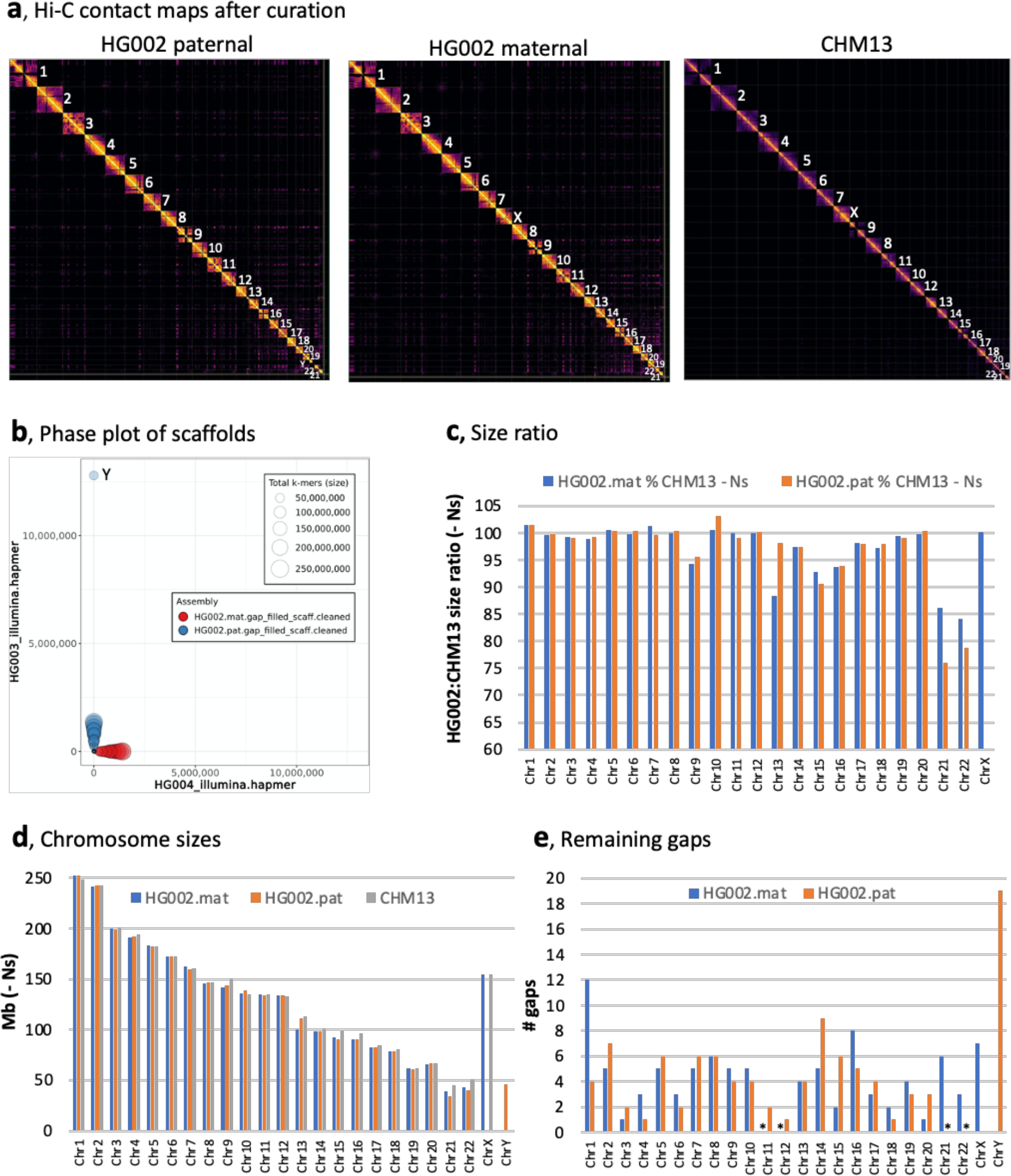
Near complete haplotype separation of scaffolds. **a**, Hi-C contact maps to the final curated HPRC-HG002 maternal and paternal assemblies in comparison to the CHM13 assembly. Values designate chromosome numbers, from largest to smallest size for each assembly. **b**, Blob plot using Illumina parental *k*-mers for the scaffolds of the HPRC-HG002 haplotypes. **c**, Percent size of HG002 diploid assembled chromosomes relative to CHM13 chromosomes, without including Ns. **d**, Absolute chromosome size values of all three assemblies compared, without including Ns. **e**, Number of remaining gaps in the chromosomes of each HG002 haplotype. *Maternal Chr 11 and 12, and paternal the remainder of Chr 21 and 22 without complete short arms have no gaps.

Despite the high levels of contiguity among the assemblies, manual curation using gEVAL alignments^37^, Bionano maps, and Hi-C interaction plots (**Extended Data Fig. 4a**) revealed a handful to several hundred contigs with missed joins, misjoins between unrelated sequences, other scaffolding errors such as inversions, translocations, and false duplications, as well as errors within contigs such as chimeric and collapsed sequences (**Supplementary Fig. 1a-d, Supplementary Table 2h**). There was no one approach, without using a highly curated reference (i.e. CrossStitch or MaSuRCA; asm1, 15, and 17), that was free of one or more scaffolding or contig errors in an automated process. For a complementary, quantitative measure of structural accuracy, we used Strand-seq data, generated by a method that selectively sequences the plus (Crick) and minus (Watson) strands of genomic DNA from cultured cells^38, 39^. Nearly all assemblies had 1-25 (average 6.5) misorientation errors (inversions or reverse complements), totalling from 1 to ∼746 Mb (**Extended Data Fig. 5a; Supplementary Table 2i**). An exception was asm14, which used Strand-seq for scaffolding. The non-Strand-seq assembly with the least misorientation errors was Trio hifiasm (asm9a,b), with only 1-2 small inversions. Over half of the assemblies had 1-9 chimeric contig errors (average 2.6), with the Trio hifiasm paternal (asm9a) assembly having the most (**Extended Data Fig. 5a; Supplementary Table 2i**). Overall, each approach avoided at least one type of error that others did not.

**Figure 4.**
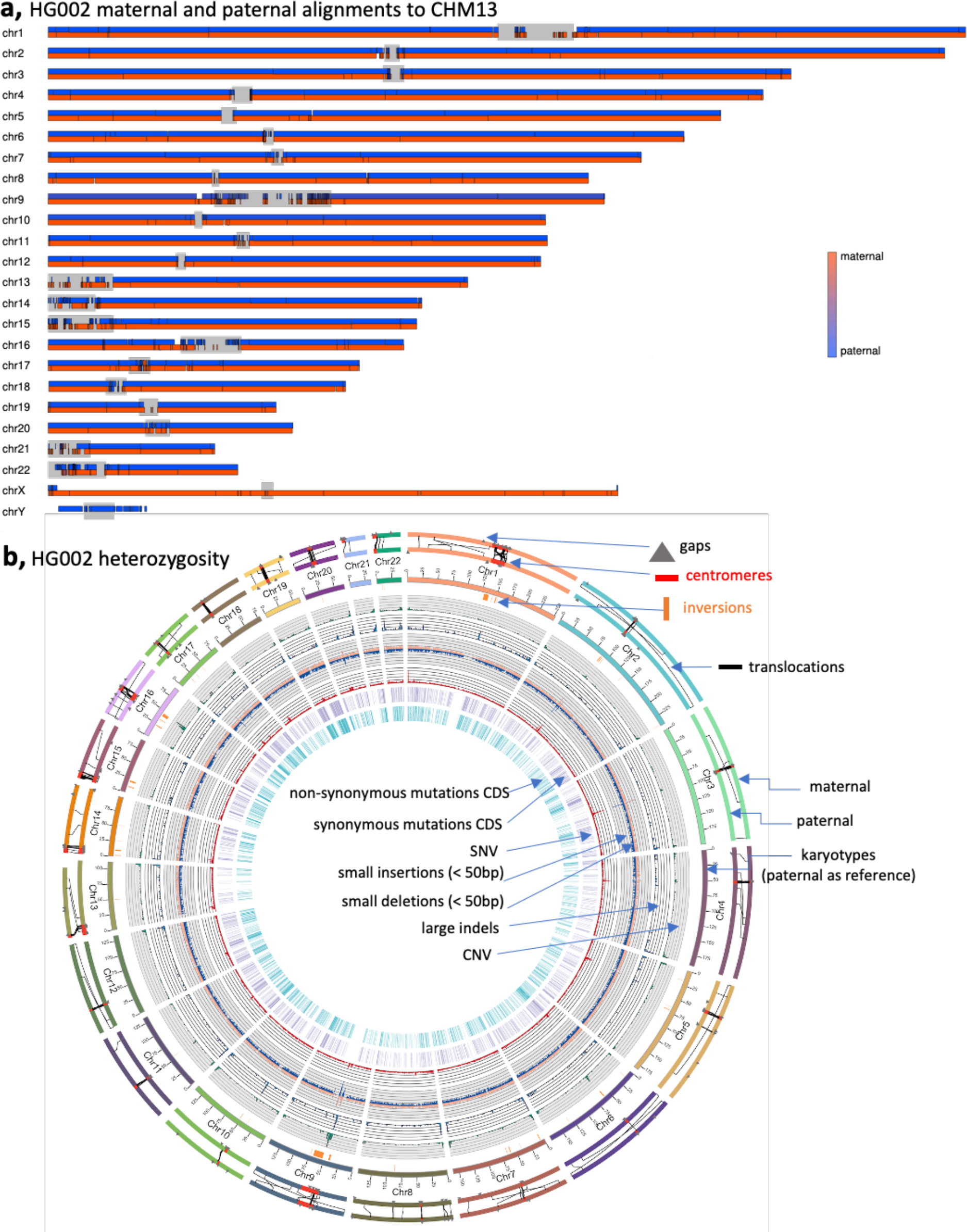
HPRC HG002 features. **a**, Chromosome (scaffold) alignments between HPRC-HG002 maternal and paternal assemblies and CHM13. Haplotype separation is nearly complete, and thus colors are solid blue (paternal) and red (maternal). Black tick marks, gaps between contigs or unaligned divergent sequence with CHM13. Alignment gaps represent structural variation or missing sequence in the HG002 assembly compared to CHM13. Grey, regions enriched with centromeric satellites. The Y chromosome reference is from GRCh38, since CHM13 does not have a Y. **b**, Circos plot of heterozygosity landscape between the two haploid assemblies of HG002. Tracks from inside out: synonymous amino acid changes; non-synonymous changes; SNV density (window size 500kb, range 0∼3.1%), small deletion and small insertion (< 50 bp) density (window size 1 Mb, range 0∼850); large indel density (≥ 50 bp, window size 1 Mb, count 0∼20), CNV density (window size 1 Mb, count 0∼77). The black line links in the outermost circles denote translocations between paternal (inner) and maternal (outer) assemblies. Orange bars, inversions (≥ 50 bp). Red bars, centromeres, Grey triangles, gaps.

**Fig. 5.**
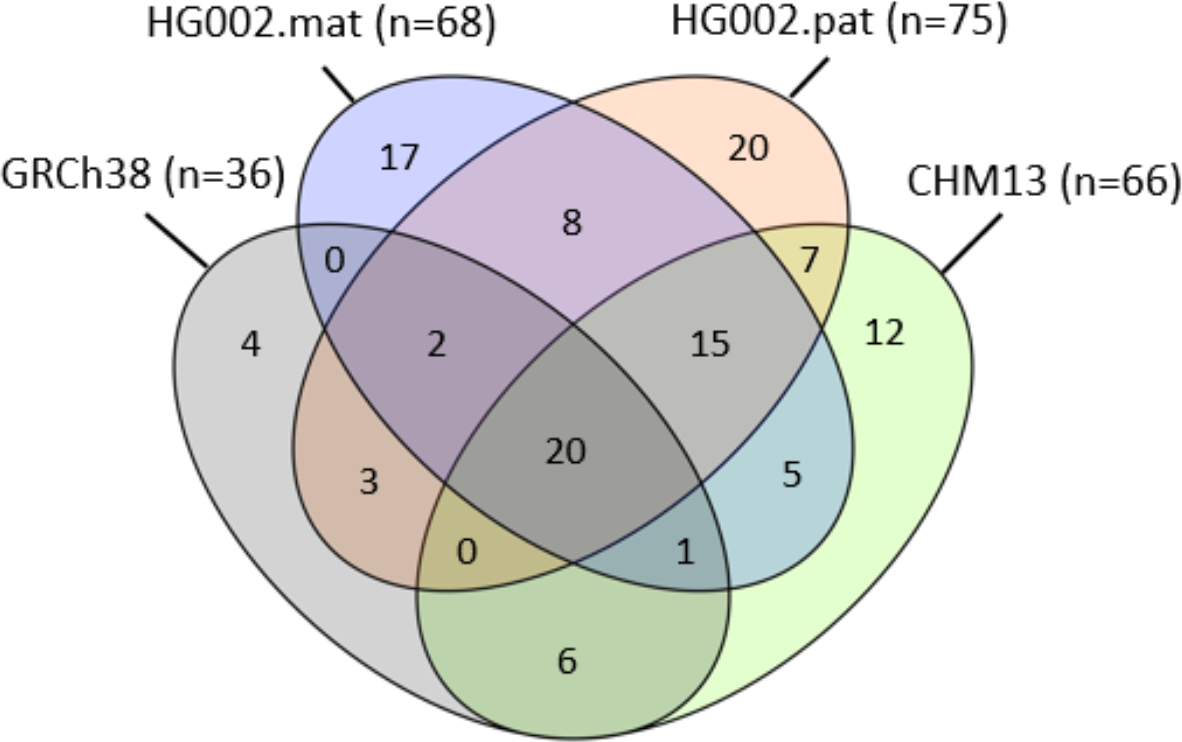
Genes unaligned and thus presumed absent in the four main reference assemblies compared. n = # of genes absent in reference assemblies. Values in the 4-way Venn diagram are the number of shared or uniquely absent genes among the 4 assemblies.

### Base call accuracy

Assembly base accuracy is critical for subsequent annotation of protein-coding genes and non-coding regulatory DNA as well as for the characterization of genetic variation, especially with complex regions that have traditionally been collapsed or excluded as a result of their repetitive content. To estimate base accuracy, we compared *k-mer* frequencies between unassembled Illumina sequencing reads and each assembly. PGAS Strand-seq (asm14a,b) generated the highest base call accuracy (QV) among scaffolded diploid assemblies while Trio hifiasm (asm9a,b) and HiCanu (asm8a) generated the highest among the contig-only diploid assemblies (QV ≥ 50, or no more than 1 base call error per 100,000 bp; **Fig. 1d; Supplementary Table 2j**). Among the merged haploid assemblies, Fly ONT.phap (asm5) performed the best, with 2 rounds of base call polishing each with ONT and HiFi reads. What these four assemblies share in common is the use of HiFi reads, either for high level read/contig filtering (asm14a,b, asm8a), polishing (asm5), and/or phasing of haplotypes (asm9a,b). Obtaining such a high degree of accuracy (QV ≥ 50) with long reads has only been a recent advance, due to the high base calling accuracy of HiFi reads^15^.

### Variant benchmarking

One of the primary goals of the HPRC is to detect and catalog genetic variation among haplotypes in the population. To determine how well each assembly detects haplotype variation, we developed a benchmarking pipeline based on variant calling. We first aligned each assembly to GRCh38, then used dipcall^38^ to call variants and compared them to a manually validated ground truth, the v4.2.1 small variant HG002 benchmark from GIAB^24^, following the Global Alliance for Genomics and Health (GA4GH) benchmarking best practices^39^. We found that all of the diploid-based assemblies had high true positive (TP) rates (aka recall or sensitivity) above 90% for single nucleotide variants (SNVs), whereas the haploid assemblies were all around 40%, due to merging of haplotypes that exclude many heterozygous variants (**Extended Data Fig. 6a; Supplementary Table 3a**). The Trio hifiasm graph-based haplotype phasing assembly (asm9) had the highest TP rate (99.47%). As expected, the values were higher (65-74%) for the haploid assemblies when ignoring genotype (GT; when only one haplotype has to match the benchmark variant), but still not as high as the diploid assemblies (**Supplementary Table 3a**). When examining variants in segmental duplications alone, the diploid assemblers performed poorly (TP dropping by 9-32%), with the exception of Trio hifiasm (94.10%) and Trio HiCanu (91.38%; which only dropped by 5 and 6% respectively; **Supplementary Table 3a**).

We next assessed the accuracy of small indels (insertions/deletions < 50 bp) between haplotypes, which are particularly problematic and highly variable due to their association with short tandem repeats (STRs). All HiFi-based diploid assemblies (∼92-98%) outperformed the haploid assemblies (∼38-59%), as well as the ONT diploid assemblies (∼52-58%; **Extended Data Fig. 6b; Supplementary Table 3b**), the latter due to the high indel error rate in ONT reads. The Trio hifiasm (asm9a,b) assembly had the highest combination of TP rates for both SNVs and small indels. The Trio hifiasm assembly was used to further improve the GIAB benchmark for SNVs, small indels, and larger structural variants (SVs; indels, inversions, and translocations) in 273 challenging, medically-relevant genes that were not well-represented in the GIAB v4.2.1 benchmark or GIAB v0.6 SV benchmark. Extensive curation by GIAB found that the Trio hifiasm assembly produced more accurate variant calls across SNVs, small indels, and SVs in these challenging regions, and the primary error type fixed was inaccurate genotypes in highly homozygous regions, particularly for indels in long homopolymers^40^. These results demonstrate that diploid assemblies are not only highly concordant but exceed existing variant benchmarks in regions resolved by mapping-based methods. They, thus, show the greatest promise for resolving more challenging regions and variants not included in current benchmarks.

### Annotation

We performed annotation for each assembly by aligning to them the human NCBI RefSeq transcriptome dataset of 78,492 transcripts from 27,225 autosomal genes, and measured mapping statistics, using GRCh38 and CHM13 assemblies as controls. Surprisingly, most of the HG002 assemblies had 100-400 genes with no transcript alignment (over 1,600 for the haploid wtdbg2 asm13 assembly; **Extended Data Fig. 7a**; **Supplementary Table 2k**). Exceptions were the Trio VGP (asm23a,b), Trio HiCanu (asm19a,b), Trio hifiasm (asm9a,b), and reference-based assemblies (asm1, 15, 17) with only ∼60-70 unaligned genes for each haplotype, twice the missing number of 36 for GRCh38 but similar to 66 genes for CHM13. Most of the contig-only assemblies had more genes (∼100-500) split between contigs than the scaffolded assemblies (**Extended Data Fig. 7a**; **Supplementary Table 2k**), consistent with scaffolding also bringing separate parts of more genes together. The Trio VGP (asm23a,b) scaffolded assembly and Trio hifiasm (asm9a,b) contig-only assembly had the fewest split genes (∼30-40) among the *de novo* assemblies, the reference-based assemblies had even fewer (1-9), and less than the GRCh38 reference (10 genes). Most assemblies had 100-700 genes (over 4,000 in the alts of asm16b and asm8b) that were less than 95% complete, except for the Trio VGP, Trio hifiasm, and reference-based assemblies with 32-89 genes (**Extended Data Fig. 7b**). For almost all assemblies, there were 200-600 genes apparently collapsed as assessed by overlapping transcript mapping, with those that used HiFi performing best (**Extended Data Fig. 7c**). Similarly, the number of genes that required frameshift error corrections were ∼1,000 for assemblies that used CLR reads (Trio VGP, asm 23a,b), ∼1,500 that used the 25kb longer but less accurate HiFi reads (asm3, asm4), ∼6,000- 16,000 (more than half of the genes) that used unpolished ONT reads, but only ∼100-200 genes with the shorter (15kb) but more accurate HiFi reads (**Extended Data Fig. 7a**; **Supplementary Table 2k**). These findings demonstrate that a critical combination of read length, base call accuracy, structural accuracy, and haplotype phasing are necessary to obtain the most complete and accurate annotation possible.

### Trio approaches yielded higher phasing accuracy

The original Trio assembly approach of binning long reads into their respective maternal and paternal haplotypes before generating contigs was originally implemented with the Canu assembler, as TrioCanu^20^, but this approach had not yet been tested in a head-to-head comparison with different assemblers and data types. Here we tested haplotype-binned reads with different contig assembly algorithms (Flye, HiCanu, HiFiasm, and Peregrine), different long read data types (HiFi, CLR, and ONT), and with trio-sorted scaffolding data types (linked reads, optical maps, and Hi-C). We found that all trio-based approaches yielded more extended phasing of the same haplotype than their non-trio counterparts. Trios that used HiFi or CLR data had the largest NG50 haplotype phase blocks (∼10-30 Mb vs < ∼0.2-5.0 Mb; **Fig. 1c**), lowest haplotype switch errors within contigs or scaffolds (∼0.01-0.02% vs 0.20-7.3%; **Extended Data Fig. 8a**), the highest number of phased bp (**Extended Data Fig. 8b**), and the most complete separation (∼99%) of paternal and maternal haplotype k-mers when using HiFi reads (**Extended Data Fig. 8c; Supplementary Table 2l,m**). Several of the trio approaches (Trio HiCanu and Trio hifiasm) yielded the least collapsed sequence (**Extended Data Fig. 9a-c; Supplementary Table 2n**). The only non-trio method that approached the phasing accuracy for maternal and paternal alleles of a trio method used Strand-seq for phasing and scaffolding (asm14; **Fig. 1c; Extended Data Fig. 8a**), but it suffered from having the highest within-scaffold errors (**Supplementary Fig. 1c**). The trio-based ONT contig assemblies had lower haplotype phase blocks (NG50s ∼3-6Mb; **Fig. 1c**), and higher haplotype switch errors (∼0.3-0.5%; **Extended Data Fig. 8a**), presumably due to their higher sequence error rates. In contrast to prior findings^9^, the VGP trio assemblies did not have the lowest haplotype false duplication rates, as assessed by either k-mers or BUSCO duplicate gene copies (**Extended Data Fig. 2d, Supplementary Table 2g**). This appears to be due to improvements in the higher read accuracy of Pacbio HiFi vs CLR, the latter used for the VGP trio assembly.

### Graph phasing assembly is most complete and accurate

The trio-based approaches fell into two principal categories: 1) those that use parental reads to haplotype bin the child’s reads before assembly (e.g. Trio VGP, Trio Flye, Trio HiCanu, Trio Peregrine); or 2) those that generate an assembly graph of the child’s genome first and then label haplotypes in the graph using the parental reads (e.g. Trio hifiasm). As presented in a complementary study conducted simultaneously^29^ and further advanced here, we found that the graph-based phasing approach generally outperformed the two-step binning trio approach when high-accuracy long reads were used to build the initial assembly graph. In particular, among the diploid assemblies, the Trio hifiasm maternal (asm9a) and paternal (asm9b) assemblies had the highest combination of high-quality metrics, including the highest QV (**Fig. 1d**), the third highest NG50 haplotype phase blocks (**Fig. 1c**; Trio VGP was the highest), among the least false duplications (**Extended Data Fig. 2d**), the fewest contigs (**Extended Data Fig. 3a**), among the lowest haplotype switch errors (**Extended Data Fig. 8a**), the highest genome completeness (**Supplementary Table 2k**), and the least collapsed repeats (**Extended Data Fig. 9a,b**). These findings indicate that a graph-based phasing assembly is more accurate and complete as the combination of the graph with haplotype information can correct errors made by either method alone. A prerequisite to highly accurate graph-based haplotype phasing is a well-resolved diploid assembly graph, as generated from high-accuracy long reads (e.g. HiFi).

### Shared properties of a pan-assembly alignment

To identify both shared and distinct features of the assemblies, we utilized a pangenomic approach, performing an all-vs-all alignment for 45 assemblies (both haplotypes each; **Extended Data Fig. 10a**), excluding the alternate contigs or unitigs of pseudohaplotype assemblies as they were highly fragmented. We annotated the alignment according to chromosomes in GRCh38 and CHM13. Pairwise Jaccard similarity analyses on the autosomes (Chr 1-22) clustered the Trio hifiasm and Trio HiCanu assemblies as more similar to each other and distinct from the other assemblies (**Fig. 2a**); at one branch higher, these trio assemblies clustered with the other trio assemblies (except Trio HiPeregrine) and with the MaSuRCA and CrossStich reference-based assemblies. The remaining super-cluster of assemblies subclustered mostly by assembly pipeline, indicating that assembly approach drives their similarities the most. More pronounced than the autosomes, the Jaccard similarity analyses on the XY sex chromosomes grouped all of the trio paternal assemblies in one cluster, with distinctions amongst themselves, relative to all of the remaining assemblies into a sister supercluster with the trio maternal assemblies (**Fig. 2b**). This finding is consistent with Chr X and part or none of Chr Y being present in both haplotypes with non-trio assemblers (**Supplementary Table 2d**). Two exceptions were the haploid Flye ONT.phap assembly (asm5) and the reference-based CrossStich hap1 assembly (asm17a), which grouped with the trio paternal assemblies and had a more complete Y chromosome (asm17a) due to using the GRCh38 Y as a reference. PCA multidimensional analyses on Euclidean distances between assemblies supported these conclusions, where in the 4th dimension the trio-based autosomes (concatenated 1 through 22) clustered by parental haplotype without the presence of the sex chromosomes (**Fig. 2c,d**). The Trio hifiasm and Trio HiCanu autosome assemblies were the most distinctly clustered by parental haplotype. Clustering on each autosome alone and then performing a machine learning algorithm (Support Vector Classifier) to find if there exists a dimension with a hyperplane that distinctly and maximally separates the trio-based maternal and paternal haplotypes, revealed such a dimension (1st to 9th, most often the 2nd), explaining 3-12% of the clustering variance (**Supplementary Table 4**). The degree of separation (i.e. PCA % variance) negatively correlated with the relative size of the centromere for each autosome (**Fig. 2e**). These findings indicate that the trio-based assemblies have the maximal separation of parental haplotypes, the centromeres contribute less to this signal, and this serves as a benchmark for further developing tools for better separation of haplotypes in non-trio assemblies.

### Combined approaches for an HPRC-HG002 reference

Based on our findings, we developed a pipeline that combines the best practices of the different approaches and used it to generate a higher-quality diploid *de novo* assembly (**Extended Data Fig. 10b**). We first removed the remaining HiFi reads with unremoved vectors (adaptors) using HiFiAdapterFilt (**Supplementary Note 2**). We then generated HiFi maternal and paternal contigs with the graph-based haplotype phasing of Trio Hifiasm v0.14.1. This updated version incorporates bug fixes we found after generating the initial HG002 assemblies, including: a) enhancing contig QV by constructing the contig golden path through high-quality portions of error corrected reads; b) resolving more segmental duplications by selecting high-occurrence seeds at the overlapping stage; and c) improving contig N50 by rescuing contained reads that break contigs on one haplotype when the read actually comes from the other haplotype^41^ (**Supplementary Fig. 2**). In addition, we titrated child and parental coverages with hifiasm and found a level (∼130x child HiFi; ∼300x parent Illumina) given the data that yielded an optimal contiguity and lowest haplotype switch error (**Supplementary Fig. 2**). We then separately scaffolded the paternal and maternal HiFi-based contigs with paternal and maternal Bionano optical maps. Conflicts between the HiFi contigs and Bionano optical maps were manually evaluated, where we accepted 5 of 15 maternal and 3 of 13 paternal joins or breaks indicated by the Bionano maps (**Supplementary Table 5a**). The majority of these conflicts (25 of 28) were in segmental duplications, in centromeres, particularly of the acrocentric chromosomes (Chr 15, 21, 22), and included natural haplotype SV differences in HG002; the remaining three were in known tandemly repeated genes (*IgK*, *IgH*, and *TSP*), where the first two are processed by programmatic structural variation associated with B lymphocytes. We then further scaffolded the paternal and maternal assemblies with haplotype-filtered (Meryl) Hi-C (OmniC) data and the Salsa 2.3 algorithm. Scaffolding with Arima Hi-C v2 data yielded similar results. We performed manual curation using Hi-C contact maps, which resulted in ∼10 breaks and ∼50 additional joins in each haplotype assembly (**Extended Data Fig. 4b**, **Supplementary Tables 2h, 5b**). Most of the breaks were at centromeres to allow satellite placement.

Next, we performed gap filling with a conservative version of the pipeline used to fill the gaps in the initial draft of the T2T CHM13 assembly^5^. ONT-UL reads were base recalled with Guppy 4.2.2, haplotype binned using trio-Canu, and then assembled into haplotype specific contigs using Flye. Draft ONT-UL contigs were polished to increase consensus accuracy. Variant calls were generated using Medaka on ONT long reads, and filtered with Merfin^42^ using kmers from Illumina short reads and then applied to increase the quality of the consensus sequence. The polished contigs were aligned to their respective haplotypes of the curated HiFi-based scaffolds from the Hi-C step above, and used to fill gaps. This resulted in 10 and 5 gaps filled in the maternal and paternal assemblies, respectively. Of these 15 gaps, 10 contained GA-rich repeats and 2 were long segmental duplications (**Supplementary Fig. 3**). The final manual curation fixed 37 items in the maternal and 60 in the paternal assemblies (**Supplementary Tables 2h and 5b**), much fewer than the hundreds of manual fixes that normally would be required (e.g. **Extended Data Fig. 4a**). A contamination screen removed multiple copies (41 maternal; 45 paternal) of endogenous human EBV viral genomes, present as single contigs, as well as a yeast contig in the paternal assembly; we did not find any non-human contamination within the human contigs and scaffolds. Approximately 98% of the remaining assembled sequence was assignable to the 22 autosomes and the X and Y sex chromosomes (**Fig. 3a**). These new assemblies were named HPRC-HG002.mat.v1.0 and HPRC-HG002.pat.v1.0 references.

These two *de novo* assemblies had the highest quality of all metrics compared to the bakeoff assemblies, including the referenced-based assemblies and the GRCh38 reference itself: the largest contig (62.9 and 81.6 Mb) and comparable scaffold (154.4 and 146.7 Mb) NG50s, close to the theoretical maximum for the scaffold values (**Fig. 1a,b**); the fewest contigs and gaps in scaffolds (**Extended Data Fig. 3a-c**); the highest QVs (∼60; **Fig. 1d**); the most complete haplotype phasing (**Fig. 3b**, **Extended Data Fig. 8a-c**) with NG50 phase blocks of 106.7 and 90.4 Mb, respectively (**Fig. 1c**, **Supplementary Table 2j**); the least collapsed repeats (18.5 and 17.6 Mb, respectively; **Extended Data Fig. 8a,b**); and among the highest annotation metrics (**Extended Data Fig. 7a-d; Supplementary Table 2m**) and the highest SNV and small indel TP rate (**Supplementary Table 3a,b**). They clustered closest with the Trio hifiasm and Trio HiCanu assemblies (**Fig. 2**). Assessing against GIAB HG002 benchmarks further, this diploid assembly produced highly accurate SNV concordance (F1 score) of 99.7% and small indel concordance of 98.6% when aligning HPRC-HG002.mat and HPRC-HG002.pat haplotypes to GRCh38. These concordance values were 0.2% and 0.8% lower than the best-performing mapping-based variant callers in a 2020 precision FDA Truth Challenge^43^. We found that 70% of the discordant SNVs fell in segmental duplications, most in the regions of complex SVs that could not be accurately benchmarked. In fact, many of these differences appeared to be more accurate in the new HPRC-HG002 assemblies than in the mapping-based benchmark or precision FDA entries. The primary limitation of the assemblies were small indels in homopolymers and in tandem repeats 51 to 200 bp long in the reference, making up 80% of all discordant indels; curation revealed that the final HPRC-HG002 assemblies had infrequent errors due to collapsing haplotypes and/or to noise in the starting HiFi reads. When benchmarking larger SVs in the new HG002 assemblies with respect to GRCh37 against the GIAB v0.6 SV benchmark, which excludes segmental duplications and centromeres^44^, TP rate was 98% (compared to 93% for asm9a,b) and precision was 89%, and most putative errors were actually just differences in representation of SVs in tandem repeats or errors in the benchmark. We found that some repetitive gene families that are known to be difficult to assemble were completely assembled in one contig, including the ∼5 Mb histocompatibility complex (MHC) containing over 220 genes (**Extended Data Fig. 10c**), where variants were > 99.99% concordant with the GIAB v4.2.1 benchmark. Overall, this high concordance between the assembly-based variants, existing benchmarks, and higher accuracy than the benchmarks, demonstrates substantial promise for phased, whole genome assemblies.

Most of these metrics, particularly for the HG002 maternal haplotype, were on par with the T2T CHM13 v1.1 assembly (**Fig. 1; Extended Data Figs. 2-3 and 7-9**), including comparable Hi-C profiles (**Fig. 3a**). We aligned the two HG002 haplotype assemblies to CHM13 (+Y from GRCh38), and found high correlations in sequence lengths (**Fig. 3c-d; Supplementary Table 6**). The HG002 chromosomes were nearly completely phased for each haplotype (**Figs. 3b** and **4a**). The unaligned sequences in the centromeres were due to much greater divergence in the centromeres; the two ends of Chr Y aligned to CHM13 Chr X, because the psuedoautosomal region (PAR) region at the ends of the HG002 Chr Y has higher identity to CHM13 Chr X than to GRCh38 Chr Y. Overall, most chromosomes (17 maternal and 17 paternal) were within 97-100% of CHM13’s length (**Fig. 3c,d**); Chr 9 was the expected size, but 10% smaller than CHM13 due to a known ∼10 Mb large satellite duplication in CHM13^5^. The biggest exceptions were the short arms of the acrocentric Chrs 13, 14, 15, 21, 22, with Chr 21, 22 being the two outliers at ∼85% of CHM13’s length for the maternal and ∼75% for the paternal haplotype (**Fig. 3d**); the short arms of these chromosomes are notoriously difficult to assemble owing to their highly repetitive structure consisting of rDNA arrays, satellite arrays, and segmental duplications^5^. Yet, the remainder of the paternal Chr 21 and 22, as well as maternal Chr 11 and 12 had no gaps, and the remaining autosomes had an average of 4 gaps (range 1-12; **Fig. 3e; Supplementary Table 6**). Most of these remaining gaps were in centromeres or acrocentric regions (**Fig. 4b**).

These findings prompted us to determine if any of the chromosomes were telomere-to-telomere complete, and thus we examined the hard to assemble regions, centromeres and telomeres. We performed diploid HiFi sequence coverage and *k-mere* analyses and found that the centromeres of 5 chromosomes (maternal 11, 12, 16 and paternal 21, 22) of 46 chromosomes have no haplotype switch errors, no collapsed repeats, and no gaps (**Extended Data Fig. 9d**; **Supplementary Figs. 4,5, Supplementary Table 7a-c**). Most of the chromosomes did not have switch errors in their centromeres, about half have some collapsed repeats, and most had gaps. We searched for canonical telomere repeats (TTAGGG), and found complete telomere q- and p-arms for 6 maternal and 10 paternal chromosomes, whereas nearly all others had one or the other arm (**Extended Data Fig. 11a-c**; **Supplementary Table 7d**). The approximately 70 unlocalized scaffolds and the several hundred remaining small unplaced scaffolds were largely centromeric satellites and telomeric repeats (**Supplementary Table 7e**). Overall, there was no chromosome that was T2T, but most are nearly complete, with few errors in centromeres or missing telomeres. These findings highlight that a mostly automatically generated, phased, and near T2T assembly is now possible, and the remaining development needed is for the centromeres and telomeres.

### Missing genes among individuals and haplotypes

The similar number of autosomal genes that did not align to both HG002 haplotypes (68 mat and 75 pat) and CHM13 (66; **Supplementary Table 2m**) led us to manually curate this annotated data set for assessing possible gene loss diversity between individuals. We found 20 genes absent from all four reference assemblies (GRCh38, HPRC-HG002.mat, HPRC-HG002.pat, and CHM13), and greater overlap of 74% (50 of 68 genes) not present in CHM13 and one or both HG002 haplotypes (**Fig. 5**). The inverse had lower overlap, with 60% (41/68) for the HG002.mat and 56% (42/75) for the HG002.pat haplotype also absent in CHM13. Similarly, the two maternal and paternal haplotypes of HG002 shared 66% (45/68) and 60% (45/75) of gene loss with each other, respectively. Conversely, CHM13 and each HG002 haplotype had 12- 20 genes absent specific to them (**Fig. 5**). However, 15 of the total HPRC-HG002.pat unaligned genes were present in one or more of the Trio paternal bakeoff assemblies, indicating that either they were missed in the HPRC-HG002.v1 reference assemblies or were false haplotype duplications in the bakeoff assemblies. False haplotype duplications are possible, given that more than half (∼67%) of the 120 genes missing among one or more of the four reference assemblies were in repetitive gene families (**Supplementary Table 8**); these include the MHC HLA immune cluster, keratin associated proteins (KRTAP), olfactory receptors (ORs), 18S and 5-8S RNA genes, several Long Intergenic Non-Protein Coding RNA (LINC) genes, and over 30 microRNA (MIR) genes. However, most genes absent in the final references were absent in all of the Trio-based assemblies, indicating real biological loss differences among individuals and haplotypes within an individual. There were about a dozen genes present in GRCh38 and the asm17 assembly that used it as a reference, but not in any of the other HG002 assemblies or CHM13, illustrating the bias of reference-based assembly methods.

### Greater diversity between haplotypes

Given the most complete human diploid assembly that we are aware of, we performed heterozygosity analysis between haplotypes, following approaches we used on a VGP trio-based marmoset assembly^45^. We noted a remarkably high amount of autosome heterozygosity (3.3% in total bp, including ∼2.6 million SNVs; ∼631K small SVs *<* 50 bp; 11.6K large SVs ≥ 50 bp; or 3,294,604 variants total; **Fig. 4b; Supplementary Table 9**). Most of the additional variation was in the newly assembled centromeres, where there were sharp peaks in SNVs, indels, inversions, and translocations (**Fig. 4b**). This could be partially explained by difficulties in detecting reliable homologies (i.e. alignments or missing sequence) in highly repetitive centromeric satellites, but which in turn can be due to higher diversity between haplotypes. When not including the centromeres, autosomal heterozygosity in total bp was ∼3-fold less (1.2%; **Supplementary Table 9**). The increased diversity in the centromeres, although expected, was not seen at this level in the marmoset trio assembly^45^. This difference could be due to marmoset assembly using higher error CLR Pacbio reads leading to more collapsing of the centromeric repeats. The alternative is that this could be due to species differences in centromere diversity, which can be determined with future assemblies using the approach developed here.

The SVs included 59 large (> 500bp) inversions, with the largest being 15 Mb on Chr 9 (**Fig. 4b; Supplementary Table 10**). Of 41 for which we had clear Watson-Crick Strand-seq alignment orientations, 30 inversions had the correct orientations, but 3 paternal and 8 maternal had the incorrect orientation (**Extended Data Fig. 5b-c**). The source of these few orientation errors appeared to be long stretches of segmental duplications on either side of the inversions, where either orientation aligns (**Extended Data Fig. 5d-f**). The SVs included 7,892 copy number variations (CNVs) between haplotypes (**Supplementary Table 9**), of which 220 were protein coding gene expansions relative to GRCh38 from 81 gene families (**Supplementary Table 11**), ∼3-fold higher than the average of 75 genes determined from less complete short-read assemblies from the 1000 genomes project^46^. Of these, 4 genes had remarkable differences of copy number or gene structure between haplotypes (**Fig. 4b**): 1) tandem arrays of “family with sequence similarity 90 member A” (*FAM90A*) members present at 32 copies in the maternal, 20 in the paternal, and 16 in GRCh38; 2) an expansion of the “nuclear pore complex interacting protein member B8” (*NPIPB8*) with 6 maternal, 10 paternal and 6 GRCh38 copies; and 3) Tre-2, Bub2p, and Cdc16p domain family member 3 (*TBC1D3*), with 11 maternal, 17 paternal and 13 GRCh38 copies; and 4) an expansion of 9 copies of the Kringle domain in lipoprotein A (*LPA*) in the paternal versus the maternal haplotype (**Supplementary Table 11**). Raw HiFi read coverage analyses of these and other examples listed did not show evidence of collapsed repeats (resolved in **Supplementary Table 11**), indicating that the haplotype differences are not assembly artifacts. The first two genes (*FAM90A, NPIPB8*) are thought to be primate specific or more rapidly evolving in primates^47, 48^; *TBC1D3* is only found in great apes, and is associated with increased cortical brain folding and expansion in humans^49^; additional copies of the Kringle domain of *LPA* has been associated with increased atherosclerosis and coronary artery disease^50^. Among the 12,736 SNVs located in CDS, 5,844 were nonsynonymous and 6,397 (50%) were synonymous, changing the amino acid sequence between haplotypes (**Fig. 4b**). Overall, a more complete human diploid genome assembly reveals a greater amount of genetic diversity than otherwise expected, and more copy number variation in genes associated with primate specific-traits.

### A look towards the future

This study allowed us to determine which current approaches yield the highest quality metrics for diploid maternal and paternal genomes of an individual. Key factors were the use of trio-parental sequence data to sort haplotype sequences in the child, a graph-based approach to resolve these haplotypes during the assembly process rather than before or after it, and combining different sequence data types and assembly tools where one approach captures information missed by another. Haplotype binning of reads before assembly (e.g. Trio HiCanu) could mispartition reads, leading to lower metrics. Haplotype phasing during a graph-based assembly (e.g. Trio hifiasm) could prevent such errors in partitioning reads.

These findings confirm and advance upon those recently reported by the VGP^9^, HGSVC^13^, and T2T^5^ assembly pipelines. The initial VGP pipeline used PacBio CLR reads, which were less accurate relative to the more recent Pacbio HiFi reads. The improved accuracy of the HiFi reads reduces the need for short-read polishing of the assemblies. More accurate long reads also allowed generation of larger contigs, reduction of collapsing repetitive sequences in the centromeres, and increased haplotype phasing accuracy^13^. For the latter, instead of FALCON-Unzip that had produced a more complete pseudohaplotype and a fragmented alternate haplotype, hifiasm, DipAsm, PGAS and CrossStich produce two comparable pseudohaplotypes. An advance adopted from the T2T approach used on CHM13 was development of a tool for automated incorporation of polished ONT assemblies for gap filling, but for both haplotypes, independently. We also made advances on the Trio assembly approach, by not only haplotype phasing the long reads and Hi-C reads, but also the Bionano optical maps. These advancements lead to near complete phased haplotypes, and near complete assembly of chromosome arms. All the components of the current pipeline developed here are available on the Galaxy computational platform, and in modular form with different steps that can be included or excluded (https://assembly.usegalaxy.eu/)51. What remains is developing diploid assembly methods that prevent the remaining collapses, gaps, and switch errors in the centromeric satellite arrays, large human satellite arrays, and short arms of the acrocentric chromosomes.

Based on the findings in this study, the HPRC has decided to use the trio graph-partitioning approach of hifiasm to generate the contigs of the first 47 individuals (94 haplotype genomes) that will contribute to the first human pangenome reference (BioProject PRJNA730822). Similarly, the trio and a non-trio version of hifiasm followed by the scaffolding approach used here for HPRC-HG002 have been adopted by other large-scale sequencing projects, such as the VGP, the Earth Biogenome Project (EBP), and the Darwin Tree of Life Project (DToL). Improvements have also been made to some of the other assembly algorithms since the versions tested here^27, 52–54^. But thus far, the trio-graph-based approach with trio-based scaffolding still yields the best combination of the highest quality metrics.

Future efforts will be necessary to develop a phasing method that does not require parental sequence data and works as well as a trio method. This will make it possible to generate equivalently good diploid reference assemblies for human and non-human organisms where parental data may not be available. Towards this end, using Hi-C or Strand-seq data for haplotype phasing are promising alternatives, as both data types contain within-chromosome haplotype information of an individual. To date, three methods have successfully used Hi-C, including FALCON Phase^21^, hifiasm (Hi-C)^53^ and pstools^55^, and another has used Strand-Seq^52^ to generate maternal and paternal phased long-read based human genome assemblies with fewer switch errors, including on HG002. As with trio binning, these approaches appear to work best when phasing is integrated with the assembly process, but further improvements are necessary to match or surpass the quality metrics seen with a parental trio-graph-based approach used here.

We used ONT-UL reads to fill in GA repeat-rich and other challenging sequence gaps in the HiFi-based contigs. As an alternative, the PacBio CLR reads that do not make it to HiFi accuracy levels contain some of the GA-rich repetitive sequences, and could be used to fill some of these gaps. The remaining few gaps per chromosome in the HG002 assemblies are mostly restricted to the hardest to assemble regions around segmental duplications, centromeres, telomeres, rDNA arrays, and other complex repeats, many with differences between haplotypes. Direct integration of ONT-UL data within the assembly graph and manual curation was necessary for finishing these regions in the T2T CHM13 assembly^5^. Thus, integration of both HiFi and ONT-UL data in a diploid assembly graph, combined with long-range phasing information from trios, Hi-C, or Strand-seq may soon enable automated T2T diploid genome assemblies. For those who wish to contribute assemblies to the human pangenome references, we encourage them to utilize our recommended processes to obtain the highest quality assemblies possible; we also encourage contribution of new methods to further improve the quality and completeness of human and other species genome assemblies.

Using currently available methods and technologies, we were able to complete over 98.5% of both HG002 haplotypes. The new biological discoveries made here demonstrate that even with a single individual, additional genetic diversity contributing to the human population can be found. Using these methods for the generation of additional diploid human genomes and creation of a human reference pangenome should enable a more complete picture of human genetic diversity, greater accuracy for precision medicine, and a greater understanding of the biology of genomes.

## Methods

### Cell lines

The GM24385 (RRID:CVCL_1C78) EBV-immortalized lymphoblastoid cell line of HG002 was obtained from the Coriell Institute for Medical Research, a National Institute for General Medical Sciences (NIGMS) cell line repository. This cell line was used to generate the Oxford Nanopore sequencing and Bionano mapping data. For the Illumina and Pacific Biosciences sequencing data, NIST Reference Material (RM) 8391 DNA was used, which was prepared from a large batch of GM24385 to control for differences arising during cell growth. For paternal (HG003) and maternal (HG004) samples, DNA was extracted from cell lines publicly available as GM24149 (RRID:CVCL_1C54) and GM24143 (RRID:CVCL_1C48), respectively, and Illumina sequencing of DNA from NIST RM 8392 (containing vials of HG002, HG003, and HG004) was used.

#### Genome sequencing

The sequence data used for this study (HG002 Data Freeze v1.0) is available in the following github link: https://github.com/human-pangenomics/HG002_Data_Freeze_v1.0. DNA samples were extracted from large homogenized growths of B-lymphoblastoid cell lines of HG002, HG003, anad HG004 from the Coriell Institute for Medical Research.

#### Illumina reads

Paired-end reads Whole genome data, TruSeq (LT) libraries, 300x PCR-free paired end 150bp + 40x, PCR-free paired end 250bp on Illumina HiSeq 2500, from GIAB^23^. HG002 was sequenced to 51.7x coverage, HG003 to 69.1x, and HG004 to 70.6x.

#### Long-molecule linked reads

10X Genomics reads: Chromium Genome Platform from 10X Genomics was sequenced to two depths: 51.7x coverage, and a deeper 84.4x coverage (300Gb) dataset. Additional data is available from BioProject: PRJNA527321. TELL-seq linked reads: These reads were made available from another study^56^.

#### PacBio reads

DNA was sheared to ∼20 kb with a Megaruptor 3, libraries were prepared with SMRTbell Express Template Prep Kit 2.0 and size-selected with SageELF to the targeted size (15 kb, 19 kb, 20 kb, or 25 kb), and sequenced on the Sequel II System with Chemistry 2.0 (15 kb or 20 kb libraries; 36x and 16x coverage, respectively), pre-2.0 Early Access Chemistry (15 kb, 19 kb, and 25 kb libraries; 24x, 14x and 11x coverage, respectively), and Sequel System with Chemistry 3.0 (15 kb libraries; 28x coverage). PacBio Continuous Long Reads (CLR): Libraries were prepared with SMRTbell Express Template Prep Kit 2.0, size selected to a target size (>30 kb), and sequenced on a Sequel II System with Chemistry 1.0 and Chemistry 2.0 to >60X fold coverage from 2 SMRT Cells.

#### ONT reads

All of the Oxford Nanopore Technologies (ONT) sequencing for HG0002 was run on PromethION and GridION sequencing instruments. The GridION uses MinION flow cells and the PromethION uses PromethION flow cells. Both flow cells employed the same ONT R9.4.1 sequencing chemistry. PromethION sequencing prepared libraries with the unsheared sequencing library prep protocol and used 28 PromethION flow cells to generate a total of 658x coverage (assuming 3.1 Gb genome size) and ∼51x coverage with 100kb+ readsShasta^26^. GridION sequencing prepared libraries with the Ultra-Long sequencing library prep protocol and used 106 MinION flow cells to generate a total of ∼52x coverage (assuming 3.1 Gb genome size) and ∼15x coverage of 100kb+ reads^57^.

#### Hi-C linked reads

Two different Hi-C data sets were made with two distinct protocols, to reach as uniform coverage across the genome as possible: Dovetail Omni-C (named Hi-C1) and Arima Genomics High Coverage Hi-C (named Hi-C2) protocols. For Hi-C1, about 100,000 cultured HG002 cells were processed for proximity ligation libraries without restriction enzymes. High coverage (69x) sequencing was done on a Nova-Seq (250bp PE). For Hi-C2, two libraries were prepared from two cell culture replicates, and sequenced with 2x150 bp and 2x250 bp Illumina reactions each. The combination of restriction enzymes represent 10 possible cut sites: ^GATC, G^ANTC, C^TNAG, T^TAA. The “^” is the cut site on the + DNA strand, and the ’N’ can be any of the 4 genomic bases.

#### Strand-seq

Strand-specific libraries were generated as previously described^13^, from 192 barcoded single-cells and sequenced on a NextSeq Illumina platform. The 192 barcoded single-cell libraries were pooled for sequencing of the HG002 sample. Raw demultiplexed fastq files from the paired-end sequencing run (80 bp read lengths) were uploaded for each single cell library. These data can be found at https://s3-us-west-2.amazonaws.com/human-pangenomics/index.html?prefix=HG002/hpp_HG002_NA24385_son_v1/Strand_seq/.

#### Optical maps

Bionano DLE1 data was collected with throughput of 1303 Gbp (Molecules >150 kb) and Read N50 of 293 kb (Molecules >150 kb) provided by Bionano Genomics and the Genome in a Bottle (GIAB) Consortium

### Genome assembly pipelines tested

The assembly bakeoff was an open public science experiment and evaluation, where researchers of the HRPC and anyone in the scientific community could contribute, with the goal of creating the highest quality *de novo* assembly possible, of one or both haplotypes, using an automated process. We did this by contacting known assembly experts, sending out announcements on consortium email list (e.g. HPRC, VGP, T2T, HGSVC, and GIAB), and announcements on HRPC associated websites (https://humanpangenome.org/hg002/; https://github.com/human-pangenomics/HG002_Data_Freeze_v1.0). We grouped the assembly pipelines tested into five categories according to whether contigs only or contigs and scaffolds were generated, and whether the contigs and/or scaffolds were haplotype phased or merged as a pseudohaplotype (**Table 1**). The assemblies were further classified by whether parental trio data was used, and whether they were reference based or *de novo* (**Table 1**). The assemblies were assigned ID numbers based on the order received by the consortium evaluation team, and in no part reflect order of assembly metric quality. All but two assemblies (asm3, asm23) that used PacBio data used the recommended downsampled HiFi SMRT cell runs from the 15 kb and 20 kb insert libraries totalling ∼34X coverage (https://github.com/human-pangenomics/HG002_Data_Freeze_v1.0#hg002-data-freeze-v10-recommended-downsampled-data-mix). Asm3 used the 19 kb, 20 kb, and 25 kb insert libraries. Asm23 used Pacbio CLR. The specific method details for each assembly pipeline, under each of the five major categories, are described below.

#### Diploid scaffold assemblies

Trio binning phasing VGP pipeline 1.6 (asm23):

This assembly was based on a modified version of the VGP trio pipeline 1.6^9^. All data types (Pacbio CLR, 10XG linked-reads, Bionano maps, and Hi-C2 reads) were haplotype-binned or filtered by haplotype. Briefly, CLR reads were binned (hapUnknownFraction=0.01) and assembled into contigs using HiCanu^15^ v1.8. NA24143 (maternal HG004) and NA24149 (paternal HG003) 250bp PE Illumina reads were used for binning. CLR coverage of the child (HG002) was 74x and 72x for the maternal and paternal haplotypes, respectively. To polish the contigs, the binned CLR reads were used for each respective haplotype with Arrow (variantCaller v2.3.3). The two haplotype contigs were then purged from each other using purge_dups v1.0^58^, conducted in the haploid mode (calcuts -d1) and only JUNK and OVLP were removed. To these contigs, Bionano molecules were aligned and assigned to the haplotype bin with higher alignment confidence. Bionano molecules aligning equally well to both parental haplotype contigs (alignment score discrepancies of less than equal to 10^-2^) were randomly split into two clusters equally and assigned to the bins. The same method of splitting the molecules was used for molecules aligning to neither of the parental assemblies (https://github.com/jlee-BNG/Bionano_Trio_binning - currently private). Binned Bionano molecules were then assembled to haploid assemblies. Cross-checking was then performed by aligning the paternal and maternal Bionano assemblies to the parental assemblies to identify regions where both parents shared the same allele, and the best allele was picked for the next round of trio binning and assemblies. 10XG and Hi-C reads were filtered for k-mers of the alternate haplotype using Meryl (https://github.com/marbl/meryl/tree/master/src/meryl) and a custom script part of the VGP trio pipeline 1.6 was used to exclude read pairs containing k-mers only found in the other haplotype. With this prepared data, three rounds of Scaffolding were then conducted on each haplotype, sequentially with the binned 10XG reads using Scaff10x v4.2, binned Bionano maps with Solve v3.4, and binned Hi-C linked reads with Salsa v2.2. The assemblies were not further polished since they already reached Q40 as judged by Merqury. Compute time was not tracked. The Source code is available at: https://github.com/VGP/vgp-assembly/tree/master/pipeline.

DipAsm contig and scaffolding pipeline (asm10):

This assembly is based on a protocol similar to DipAsm reported in^27^. Pacbio HiFi reads were first assembled into unphased contigs using Peregrine. Contigs were grouped and ordered into scaffolds with Hi-C2 data. The HiFi reads were then mapped to scaffolds using minimap2 and heterozygous SNPs called using DeepVariant^59^. The heterozygous SNP calls were phased with both HiFi and Hi-C2 data using HapCUT2^60^ and Whatshap^61^. The reads were then partitioned based on their phase using a custom script. The partitioned reads were re-assembled into phased contigs using Peregrine. The contigs were then ordered and joined together with 100 Ns to produce phased scaffolds. Compute time was not tracked. The source code is available at: https://github.com/shilpagarg/DipAsm

Dovetail DipAsm variant pipeline (asm2 and asm22):

The Dovetail pipeline used is a variation of the DipAsm pipeline described in^27^. The main difference is that DipAsm used HiFi reads for SNP calling with DeepVariant and the Dovetail protocol used Omni-C reads (Hi-C1) for SNP calling with Freebayes. In particular, Pacbio HiFi reads were assembled into contigs using the Peregrine assembler with default parameters. These contigs were joined into chromosome-scale scaffolds using Dovetail Omni-C data and either HiRise (Dovetail Genomics; asm2) or Salsa2^16^ (asm22) scaffolders. Omni-C reads were then aligned to scaffolds and haplotype SNPs were called using FreeBayes. These SNPs were then phased with HapCUT2 and Omni-C long-range links to obtain chromosome-scale phased blocks. These phased SNPs were used to partition HiFi and Omni-C reads into two haplotypes. Reads for which the partitioning could not be done ambiguously were assigned to both haplotypes. Phased HiFi reads for each haplotype were assembled again with Peregrine and scaffolded with haplotype-specific Omni-C reads to obtain chromosome-scale phased scaffolds. Compute time was not tracked; all the tools were run on AWS EC2 with c5d.9xlarge instance type. Source code for HiRise is proprietary. Source for SALSA2 is available at: https://github.com/marbl/SALSA/commit/974589f3302b773dcf0f20c3332fe9daf009fb93

PGAS pipeline (asm14):

The recent Phased Genome Assembly using Strand-seq (PGAS) diploid genome assembly pipeline is described in^52^. First, a non-haplotype resolved (“squashed”) contig assembly was generated from PacBio HiFi reads using Peregrine v0.1.5.5 (github.com/cschin/Peregrine). Illumina short reads from the Strand-seq data^62^ were aligned against this squashed assembly to identify contigs that most likely originate from the same chromosome based on similar Watson/Crick strand inheritance patterns^63^. This information was then used to cluster the contigs into roughly chromosome-scale clusters, which helps avoid chimeric chromosome assemblies, allows for parallelization of the assembly pipeline, and facilitates phasing. Next, heterozygous single nucleotide variants were identified based on long-read alignments against the clustered assembly with DeepVariant v0.9.0. To obtain chromosome-scale haplotypes, integrative phasing with WhatsHap^61^ was performed, combining local dense phase information derived from long reads with global sparse phase information inferred from Strand-seq alignments. Next phased heterozygous SNVs were used to assign each HiFi read to its corresponding haplotype (“haplo-tagging”) or remain in the fraction of haplotype-unassigned reads. The haplotags were used to split the HiFi reads into two haploid read sets, which, together with the haplotype-unassigned reads, were the input to assemble two haplotype contig sequences per chromosome-scale cluster with Peregrine v0.1.5.5. After polishing the contigs for two rounds with Racon v1.4.10^64^ and the haploid long-read datasets, the per chromosome cluster assemblies were merged to create a genome-scale diploid assembly. The final round of scaffolding of each haplotype was performed with the short reads from the Strand-seq data, on HiFi contigs with a minimum size of 500 kb. This size thresholding was necessary since the contig order can only be inferred from strand-state changes resulting from sister chromatid exchanges (SCE; a process during DNA replication where two sister chromatids break, rejoin, and physically exchange regions of the parental strands). SCEs are low-frequency events that are thus less likely to produce a traceable signal with decreasing contig size. The complete assembly pipeline run required less than 2000 CPU hours on a 3-node cluster (3 x 36C, 1.4 TB RAM) with a peak RAM usage of around 600 GB (squashed Peregrine assembly). Source code is available at: github.com/ptrebert/project-diploid-assembly; pipeline parameter version 8

CrossStitch (asm17):

The assembly was produced using CrossStitch, a reference-based pipeline for diploid genome assembly. SNPs and small indels were called with respect to GRCh38 for HG002 from alignments of unbinned 30x PacBio HiFi reads. Variant calling was performed on this BAM using DeepVariant v0.9^59^ and the PacBio model. A full set of commands and parameters are available on the PacBio case study: https://github.com/google/deepvariant/blob/r0.9/docs/deepvariant-pacbio-model-case-study.md. Larger structural variants (SV) were called by running Sniffles v1.0.11 (with parameters -s 10 -l 10 --min_homo_af 0.7) on minimap2 v2.17 alignments of the HiFi reads and refining these calls with Iris v1.0.1 (https://github.com/mkirsche/Iris)65. Then, the SNVs and small indel variants (less than 30 bp) called from DeepVariant were phased using HapCUT2 v1.1 on the ONT + Hi-C alignments, and these phase blocks were used to assign a haplotype to each HiFi read. SV phasing was performed by observing the reads supporting each heterozygous SV call and assigning the variant to the haplotype which the majority of the reads came from. Finally, vcf2diploid (https://github.com/abyzovlab/vcf2diploid) from the AlleleSeq algorithm^66^ was u^66^d to incorporate small variant and SV calls into a template consisting of the GRCh38 reference genome sequence, producing the final assembly for HG002. The end-to-end assembly took less than two days on a high memory machine at JHU using at most 40 cores at a time. Peak RAM utilization was <100GB Source code is available at: https://github.com/schatzlab/crossstitch (commit e49527b)

#### Diploid contig assemblies

Trio binning flye ONT pipeline (asm6 and asm7):

Following a trio based assembly approach^20^, Parental Illumina 21-mers were counted in the child, maternal, and paternal read sets (full sets, not subset coverage recommendations). Haplotype-specific mers were created using Merqury v1.0^67^ and Meryl v1.0 (https://github.com/marbl/meryl) with the command: hapmers.sh.sh mat.k21.meryl pat.k21.meryl child.k21.meryl. These short reads were then used to bin ONT standard long (asm6) or ultralong > 100kb (asm7) reads into their maternal and paternal specific haplotypes. The ONT recommended subset reads were then assigned using splitHaplotigs in Canu v2.0^15^ with the command: splitHaplotype -cl 1000 -memory 32 -threads 28 -R HG002_ucsc_ONT_lt100kb.fastq.gz \

-R HG002_giab_ULfastqs_guppy3.2.4.fastq.gz \

-H ./0-kmers/haplotype-DAD.meryl 6 ./haplotype-DAD.fasta.gz \

-H ./0-kmers/haplotype-MOM.meryl 6 ./haplotype-MOM.fasta.gz \

-A ./haplotype-unknown.fasta.gz.

Flye v2.7-b1585^28^ was then run on the binned reads to generate maternal and paternal contigs with the command fly --threads 128 --min-overlap 10000 --asm-coverage 40 -out_dir <MOM/DAD> --genome-size 3.1g --nano-raw haplotype-<DAD/MOM>.fasta.gz. Flye sometimes inserts gaps when it isn’t certain of a repeat sequence, and thus some contigs appear as scaffolds. But the assembly is still contig-level. No base-level polishing (with short or long reads) was conducted on the assembly. The ONT standard Flye runs took approximately 1200 CPU hrs (20 wall clock) and 500 Gb of memory. The ONT ultralong assemblies took approximately 3000 CPU hrs (60 wall clock) and 800 Gb of memory. Source codes for Canu, Mercury, and Flye are available at: https://github.com/marbl/canu; https://github.com/marbl/merqury; https://github.com/fenderglass/Flye

Trio binning HiCanu contig pipeline (asm19):

Following a trio based assembly approach^20^, parental Illumina 21-mers were counted in the child, maternal, and paternal read sets (full sets, not subset coverage recommendations). Haplotype-specific mers were created using Merqury v1.0^67^ and Meryl v1.0 (https://github.com/marbl/meryl) with the command: hapmers.sh mat.k21.meryl pat.k21.meryl child.k21.meryl. The HiFi recommended 34x subset reads were then assigned to using splitHaplotigs in Canu v2.0^15^ with the command: splitHaplotype -cl 1000 -memory 32 -threads 28 -R m64012_190920_173625.fastq.gz -R m64012_190921_234837.fastq.gz -R m64011_190830_220126.Q20.fastq.gz -R m64011_190901_095311.Q20.fastq.gz -H ./0-kmers/haplotype- DAD.meryl 6 ./haplotype-DAD.fasta.gz -H ./0-kmers/haplotype-MOM.meryl 6 ./haplotype-MOM.fasta.gz -A ./haplotype-unknown.fasta.gz. Any reads which were unclassified were randomly divided into two bins. The resulting maternal and paternal read sets were independently assembled with HiCanu^15^ v2.0 with the commands:

canu -p ’asm’ ’gridOptions=--time=4:00:00 --partition=quick,norm’ ’gridOptionsCns=--time=30:00:00 -- partition=norm ’ ’genomeSize=3.1g’ ’gfaThreads=48’ ’batOptions= -eg 0.01 -sb 0.01 -dg 6 -db 6 -dr 1 -ca 50 -cp 5’ -pacbio-hifi haplotype-[DAD|MOM].fasta.gz haplotype-unknown-batch[1|2].fastq.gz. Source code: https://github.com/marbl/canu, https://github.com/marbl/merqury

Publication citation: https://doi.org/10.1101/2020.03.14.992248

All runs used the “quick” partition of the NIH Biowulf cluster (https://hpc.nih.gov). HiCanu required approximately 1400 CPU hours per haplotype (19 wall clock hours) and no single job required more than 64 GB of memory.

Trio binning Peregrine contig pipeline (asm20):

The same binned reads as for asm19 were used for this assembly. The reads were assembled with Peregrine v0.1.5.3+0.gd1eeebc.dirty with the command yes yes | python3 Peregrine/bin/pg_run.py asm \ input.list 24 24 24 24 24 24 24 24 24 \

--with-consensus --shimmer-r 3 --best_n_ovlp 8 \

--output ./

with the input.list specifying appropriate haplotype input reads. Compute time was not tracked. Source code: https://github.com/cschin/Peregrine, https://github.com/marbl/merqury

Trio phasing Hifiasm contig pipeline (asm9):

Hifiasm finds alignments between HiFi reads and corrects sequencing errors observed in alignments^29^. It labels a corrected read with its inferred parental origin using parent-specific 31-mers counted from parental short reads. HiFi reads in long homozygous regions do not harbor parent-specific 31-mers and are thus unlabeled. Hifiasm then builds a string graph from read overlaps that carries read labels. It traces paternal and maternal reads in the graph to generate paternal and maternal contigs, respectively. We collected paternal 31-mers from short reads with “yak count -b37 -o sr-pat.yak sr-pat.fq.gz” (and similarly for maternal) and assembled HiFi reads with “hifiasm -1 sr-pat.yak -2 sr-mat.yak hifi-reads.fq.gz”. The assembly took 305 CPU hours. Source code is available at: https://github.com/chhylp123/hifiasm/releases/tag/v0.3.

### DipAsm contig pipeline (asm11)

The assembly pipeline mimics the DipAsm steps explained for asm10. The pipeline takes as input HiFi and Hi-C datasets and outputs the phased contigs. Initially, the pipeline produces unphased contigs using Peregrine and then these unphased contigs are scaffolded to produce chromosome-scale sequences using HiRise. Afterwards, the heterozygous SNPs are called and are phased using HiFi and Hi-C data. These phased SNPs are informative sites to partition HiFi reads to haplotypes on the chromosome level. The phased reads are then assembled using Peregrine to produce phased contigs.

### Peregrine contig pipeline (asm3 and asm4)

The Peregrine assembler^32^ was used to generate contigs on the HiFi reads, using either the full coverage sequence (asm3) consisting of 19Kb, 20kb, and longer 25Kb read libraries or downsampled to 34X and shorter 15 kb reads (asm 4). A module was written to separate likely true variant sites from differences between reads caused by sequencing errors. This was done by using the overlap data from the peregrine assembler overlapping modules with additional alignment analysis. The variants of the read overlapped data were analyzed to get a subset of variants that should belong to the same haplotypes. The reads with the same set of variants were considered to be haplotype consistent and the overlaps between those haplotype consistent reads were considered for constructing the contig assembly. Overlaps between different haplotypes or different repeats from the analysis results were ignored. It is expected that the generated contigs are from single haplotypes in those regions, which have enough heterozygous variants. Compute time was not tracked. Source code is available at: https://github.com/cschin/Peregrine_dev/commit/93d416707edf257c4bcb29b9693c3fda25d97a29

### FALCON-Unzip contig pipeline (asm16)

FALCON-Unzip^19^ version 2.2.4-py37hed50d52_0 was run on reads from 4 SMRT Cells from two HiFi libraries (15 kb and 20 kb, 34x coverage total) reads with “input_type = preads, length_cutoff_pr = 8000, ovlp_daligner_option = -k24 -h1024 -e.98 -l1500 -s100, ovlp_HPCdaligner_option = -v -B128 -M24, ovlp_DBsplit_option = -s400, overlap_filtering_setting = --max-diff 200 --max-cov 200 --min-cov 2 --n- core 24 --min-idt 98 --ignore-indels” for the initial contig assembly and default parameters for unzipping haplotypes. The assembly took 2,540 CPU-core hours on nodes with Intel Xeon processor E5-2600 v4. Source code is available at: https://anaconda.org/bioconda/pb-falcon/2.2.4/download/linux-64/pb-falcon-2.2.4-py37hed50d52_0.tar.bz2

### HiCanu purge dups contig phasing pipeline (asm8)

HiCanu^15^ v2.0 with the command canu -p ’asm’ ’gridOptions=--time=4:00:00 --partition=quick,norm’ ’gridOptionsCns=--time=30:00:00 --partition=norm ’ ’genomeSize=3.1g’ ’gfaThreads=48’ -pacbio-hifi m64012_190920_173625.fastq.gz m64012_190921_234837.fastq.gz m64011_190830_220126.Q20.fastq.gz m64011_190901_095311.Q20.fastq.gz Purge_dups^58^ was used to remove alternate haplotypes, GitHub commit b5ce3276773608c7fb4978a24ab29fdd0d65f1b5, with the thresholds of 5 7 11 30 22 42. Purge_dups introduces gaps near the purged sequenced regions of the contigs, and thus some contigs appear as scaffolds.

But the assembly is still contig-level. HiCanu required approximately 1800 CPU hours and no single job required more than 64 GB of memory (22 wall clock hours). Purge dups required 40 CPU hours and <1GB of memory.

### Haploid scaffold assemblies

Flye ONT pipeline (asm5):

Flye v2.7b-b1579^28^ was used to assemble (downsampled) ONT reads into contigs, using the default parameters with extra “--asm-coverage 50 --min-overlap 10000” options. Two iterations of the Flye polishing module were applied using ONT reads, followed by two polishing iterations using HiFi reads. Finally, Flye graph-based scaffolding module was run on the polished contigs, which generated 54 scaffold connections and slightly improved the assembly contiguity. Assembly took ∼5000 CPU hours and polishing (ONT + HiFi) took ∼3000 CPU hours. Peak RAM usage was ∼900 Gb. The pipeline was run on a single computational node with two Intel Xeon 8164 CPUs, with 26 cores each and 1.5 TB of RAM. Source code is available at: https://github.com/fenderglass/Flye/. Commit id: ec206f8

Shasta ONT + HiC (asm18 and asm21):

The Shasta assembler^26^ was used to assemble ONT reads into contigs. The contigs were polished using PEPPER (https://github.com/kishwarshafin/pepper) which also uses only the ONT reads. The contigs were scaffolded with Omni Hi-C (Hi-C1) using HiRise (asm18) or Salsa v2.0 (asm21). Compute time was not tracked. Source code is available at: https://github.com/chanzuckerberg/shasta

Flye and MaSuRCA (asm15):

A subset of downsampled ONT UL data that contained ∼38x genome coverage of 120kb reads or longer was used. The ONT reads were assembled into contigs using the Flye assembler^28^ v2.5. The contigs were polished with downsampled 30x coverage of PacBio HiFi 15kb and 20kb reads, using POLCA, a tool distributed with the Maryland Super-Read Celera Assembler (MaSuRCA)^30^. To scaffold and assign the assembled contigs to chromosomes, a reference-based scaffolding method was used embodied in the chromosome_scaffolder script included in MaSuRCA. GRCh38.p12 was used as a reference (without the ALT scaffolds) for scaffolding, with chromosome_scaffolder option enabled, which allows it to fill in the gaps in scaffolds, where possible, with GRCh38 sequence, in lowercase letters. The final assembly was named JHU_HG002_v0.1. Compute time was not tracked. Source code is available at: https://github.com/alekseyzimin/masurca

MaSuRCA (asm1):

MaSuRCA v3.3.1^30^ with default parameters was run on the combined Illumina, ONT, and PacBio HiFi data to obtain a set of contigs designated the Ash1 v0.5 assembly. The ONT and PacBio data were an earlier release, from 2018, and the read lengths were shorter than the later release used by most other methods in this evaluation. After initial scaffolding, MaSuRCA was used to remove redundant haplotype-variant scaffolds by aligning the assembly to itself and looking for scaffolds that were completely covered by another larger scaffold and that were >97% identical to the larger scaffold. To scaffold and assign the assembled contigs to chromosomes, we used a reference-based scaffolding method embodied in the chromosome_scaffolder script included in MaSuRCA. We used the GRCh38.p12 as the reference (without the ALT scaffolds), and we enabled an option in chromosome_scaffolder that allows it to fill in gaps with GRCh38 sequence, using lowercase letters. Finally, we examined SNVs reported at high frequency in an Ashkenazi population from the Genome Aggregation Database (gnomAD). GnomAD v3.0 contains SNV calls from short-read whole-genome data from 1,662 Ashkenazi individuals. At 273,866 heterozygous SNV sites where HG002 contained the Ashkenazi major allele and where our assembly used a minor allele, we replaced the allele in Ash1 with the Ashkenazi major allele. A publication of the final curated asm1 assembly has been made^54^. Compute time was not tracked. Source code is available at: https://github.com/alekseyzimin/masurca

### Haploid contig assemblies

wtdbg2 (asm13)

The standard wtdbg2 assembly pipeline^33^ was applied on HiFi-reads. Parameters ’-k 23 -p 0 -S 0.8 --no-read-clip --aln-dovetail -1’ were customized to improve the contiguity. Source code is available at: https://github.com/ruanjue/wtdbg2 commit:d6667e78bbde00232ff25d3b6f16964cc7639378

#!/bin/bash

wtdbg2 -k 23 -p 0 -AS 4 -s 0.8 -g 3g -t 96 --no-read-clip --aln-dovetail -1 -fo dbg -i

../rawdata/SRR10382244.

fasta -i ../rawdata/SRR10382245.fasta -i ../rawdata/SRR10382248.fasta -i ../rawdata/SRR10382249.fasta

wtpoa-cns -t 96 -i dbg.ctg.lay.gz -fo dbg.raw.fa

minimap2 -I64G -ax asm20 -t96 -r2k dbg.raw.fa ../rawdata/SRR10382244.fasta

../rawdata/SRR10382245.fasta

../rawdata/SRR10382248.fasta ../rawdata/SRR10382249.fasta | samtools sort -m 2g -@96 -o dbg.bam samtools view -F0x900 dbg.bam | wtpoa-cns -t 96 -d dbg.raw.fa -i - -fo dbg.cns.fa

Publication citation: 10.1101/530972 Compute time.

wtdbg2: real 20349.731 sec, user 1178897.390 sec, sys 18351.090 sec, maxrss 194403704.0 kB, maxvsize 209814736.0 kB

wtpoa-cns(1): real 3350.517 sec, user 260551.730 sec, sys 1040.200 sec, maxrss 9978492.0 kB, maxvsize 15839032.0 kB

wtpoa-cns(2):real 2181.084 sec, user 149528.810 sec, sys 815.380 sec, maxrss 11134244.0 kB, maxvsize 16012012.0 kB

others: unknown

NECAT Feng Luo group (asm12):

We used the NECAT assembler^31^ to assemble ONT reads of HG002, which contained about 53x coverage excluding ONT UL reads. The command “necat.pl config cfg” was first used to generate the parameter file ’cfg’. The default values in ’cfg’ were replaced with the following parameters: “GENOME_SIZE=3000000000, THREADS=64, PREP_OUTPUT_COVERAGE=40, OVLP_FAST_OPTIONS=-n 500 -z 20 -b 2000 -e 0.5 -j 0 -u 1 -a 1000”, OVLP_SENSITIVE_OPTIONS=- n 500 -z 10 -e 0.5 -j 0 -u 1 -a 1000, CNS_FAST_OPTIONS=-a 2000 -x 4 -y 12 -l 1000 -e 0.5 -p 0.8 -u 0, CNS_SENSITIVE_OPTIONS=-a 2000 -x 4 -y 12 -l 1000 -e 0.5 -p 0.8 -u 0, TRIM_OVLP_OPTIONS=-n 100 -z 10 -b 2000 -e 0.5 -j 1 -u 1 -a 400, ASM_OVLP_OPTIONS=-n 100 -z 10 -b 2000 -e 0.5 -j 1 -u 0 –a 400, CNS_OUTPUT_COVERAGE=40”. The command “necat.pl bridge cfg” was run to generate the final contigs. It took ∼12555 CPU hours (error correction 2500 h, assembling 8123 h, bridging 1216 h, polishing 716 h) on a 4-core 24-thread Intel(R) Xeon(R) 2.4 GHz CPU (CPU E7-8894[v4]) machine with 3 TB of RAM. Source code is available at: https://github.com/xiaochuanle/NECAT. Commit id: 47c6c23.

### HPRC-HG002 references

HiFi reads with adaptors were removed with HiFiAdapterFilt (https://github.com/sheinasim/HiFiAdapterFilt). Maternal and paternal contigs were generated from 130X coverage of the remaining HiFi reads using hifiasm v0.14.1 in trio mode. The maternal and paternal HiFi contigs were then separately scaffolded with paternal and maternal Trio-Bionano Solve v1.6 maps, respectively. Conflicts between the Bionano maps and Pacbio HiFi-based contigs were manually reviewed by three experts and decisions were made to accept or reject the cuts proposed by Bionano Solve. Further scaffolding of the paternal and maternal scaffolds was done with Salsa v2.3 and OmniC Hi-C reads excluding reads from the other haplotype with Meryl; that is, paired reads with *k*-mers only seen in one parent were removed prior to mapping and scaffolding. The resulting scaffolds were manually curated to ensure the proper order and orientation of contigs within the scaffolds, leading to additional joins and breaks. In parallel, ONT-UL reads were haplotype-binned with Canu v2.1 using Illumina short reads of the parents, and then were assembled into their respective maternal and paternal contigs using Flye v2.8.3- b1695 with --min-overlap 10000. Bases were recalled with Guppy v4.2.2 prior to the assembly. The contigs were polished calling variants with Medaka (https://github.com/nanoporetech/medaka). The variants were filtered with Merfin using k-mers derived from Illumina short reads. Bcftools was used to apply the variants. The resulting ONT-UL assembly was used to patch gaps of the scaffolded HiFi-based assembly, using custom scripts (https://github.com/gf777/misc/tree/master/HPRC%20HG002/for_filling). Finally, a decontamination and an additional round of manual curation were conducted.

### Trio binning with optical mapping

Using the Bionano Direct Label and Stain (DLS) chemistry and the Saphyr machine, high coverage of Bionano optical maps were generated of the HG002 (son), HG003 (father), and HG004 (mother) trio of samples and each assembled into diploid assemblies (**Supplementary Table 12**) with Bionano Solve 3.6. To separate the paternal and maternal alleles in the child assembly, the child molecules were aligned to the father and mother assemblies and assigned to the bin with higher alignment confidence. Molecules that aligned equally well (alignment score difference ≼ 10^-2^) to the parents were split into two clusters equally and assigned to the bins. Similarly, molecules that aligned to neither of the parents were split into two clusters equally and assigned to the bins. Since this method utilizes the unique SV sites in the parents’ diploid assemblies to bin the molecules, it does not distinguish molecules for regions where the parents have the same structural variants. Without special adjustment for the sex chromosome, this method does not eliminate the assembly of Chr X in the paternal assembly but the contigs are much shorter due to the missing proband molecules of regions where the father has unique SVs.

To further improve the separation of the parental alleles, a cross-checking step is performed. The binned paternal and maternal assemblies are aligned to both the father and mother assemblies (cross_check_alignment.py with RefAligner 11741 and optArguments_customized_for_diploid_reference.xml). Using these alignments, regions where the parents share an allele but are homozygous in one and heterozygous in the other are identified. For example, in regions where the father has allele AA and the mother has allele AB, the proband’s B allele would be from the maternal and A allele would be from the paternal side. Unless there are nearby SVs, molecules with allele A in the child can align equally to both parents, where the maternal assembly will then also include allele A, but with cross checking, the correct allele (allele B) is identified and the wrong allele (allele A) gets eliminated through breaking it in the maternal assembly. A total of 54 regions of such characteristics were identified and broken in the paternal assembly, and 45 were identified and broken in the maternal assembly (haplotype_segregation_cross_check_rscript.R, haplotype_segregation_cut_step.py). Breaking at these regions allowed further separation of alleles in the next round of trio binning, using the binned and cross-checked assemblies as anchors. For the trio-binning after cross-checking, 586 Gbp of the probe molecules were binned to the paternal haplotype and 615 Gbp binned to the maternal haplotype. These binned molecules were then assembled into the paternal haploid assembly (2.98 Gbp with N50 of 79.22 Mbp) and maternal haploid assembly (2.96 G with N50 of 66.60 Mbp) using Bionano Solve 3.6.

**Supplementary Table 12.** Molecules and Bionano map assembly metrics of HG002, HG003 and HG004. All molecules (original and binned) and assemblies can be found in https://bionanowithcentrexit-my.sharepoint.com/:f:/g/personal/jlee_bionanogenomics_com/Eq9gz0in15hKnC5CMrhUYkwBmjY0uiC_gPhu5QDR82AThw?e=u5Eyzd. All trio binning and cross-checking scripts can be found in https://github.com/andypang0714/Bionano_Trio_binning v2.0.

|  | HG002 | HG003 | HG004 |
| --- | --- | --- | --- |
| Total DNA ($\geq 150$ kbp) | 1303 Gbp | 1332 Gbp | 1304 Gbp |
| N50 ($\geq 150$ kbp) | 293 kbp | 298 kbp | 348 kbp |
| Diploid genome map length | 5.95 Gbp | 5.93 Gbp | 5.92 Gbp |
| Diploid map N50 | 71.07 Mbp | 69.94 Mbp | 81.51 Mbp |

### Evaluation methods

#### Contamination, manual curation

Curation was conducted as described in the VGP^9^. In brief, for Contamination identification, a succession of searches was used to identify potential contaminants in the generated assemblies. This included: 1) A megaBLAST98 search against a database of common contaminants (ftp://ftp.ncbi.nlm.nih.gov/pub/kitts/contam_in_euks.fa.gz); 2) A vecscreen (https://www.ncbi.nlm.nih.gov/tools/vecscreen/) search against a database of adaptor sequences (ftp://ftp.ncbi.nlm.nih.gov/pub/kitts/adaptors_for_screening_euks.fa). The mitochondrial genome was identified by a megaBLAST search against a database of known organelle genomes requiring (ftp://ftp.ncbi.nlm.nih.gov/blast/db/FASTA/mito.nt.gz). Organelle matches embedded in nuclear sequences that were found to be NuMTs. For structural error identification, for each assembly, all sequence data (CLR, HiFi, ONT, and optical maps) were aligned and analysed in gEVAL95 (https://vgp-geval.sanger.ac.uk/index.html). Separately, Hi-C data were mapped to the primary assembly and visualized using HiGlass97. These alignments were then used by curators to identify mis-joins, missed joins and other anomalies. We identified sex chromosomes based on half coverage alignments to sex chromosomes in GRCh38.

#### Continuity metrics

Assembly continuity statistics were collected using asm_stats.sh from the VGP pipeline [https://github.com/VGP/vgp-assembly/tree/master/pipeline/stats,9, using genome size of 3 Gb for calculating NG50 values. All N-bases were considered as gaps.

#### Completeness, phasing, and base call accuracies

We collected 21-mers from Illumina reads of HG002 (250bp paired-end) and the parental genomes (HG003 and HG004) using Meryl^67^, and used Merqury^67^ to calculate QV, completeness and phasing statistics. Like continuity metrics, phase block NG50 was obtained using a genome size of 3Gb. False duplications were post-calculated using false_duplications.sh in Merqury and spectra-cn histogram files for each haploid- representation of the assemblies.

#### Collapsed analyses

We calculated collapsed and expandable sequences using previously described methods^68^. In brief we aligned downsampled HiFi reads from HG002 independently to each assembly of HG002 and defined collapsed bases as regions in the assembly with greater than expected coverage (mean + at least three standard deviations) that were at least 15kb in length. We performed analyses with common repeats collapses included and excluded, defining common repeats collapses as sequences that were over 75% common repeat elements as identified by RepeatMasker (v4.1.0) and TRF (v4.09). This filter removed many collapses corresponding to alpha satellite and human satellite to get a better estimate of collapsed segmental duplications. Further, we defined expandable Mbp as the estimate of how much sequence would be in the collapsed regions had they been correctly assembled. We estimated this by multiplying the length of each collapse against the read depth divided by the average genome coverage. Code used for this analysis is available at https://github.com/mrvollger/SDA (commit 23fa175).

#### Strand-seq analyses

In order to evaluate structural accuracy of each assembly, we first aligned Strand-seq data from HG002 to each assembly using BWA-MEM (version 0.7.15-r1140)^69^ with the default parameters. Subsequently, all secondary and supplementary alignments were removed using SAMtools (version 1.9)^70^ and duplicate reads were marked using Sambamba (version 0.6.8)^71^. Duplicated reads and reads with mapping quality less than 10 were removed prior to subsequent Strand-seq data analysis. To evaluate structural and directional contiguity of each assembly, we used R package SaaRclust^52^ with the following parameters: bin.size = 200000; step.size = 200000; prob.th=0.25; bin.method = ’fixed’; min.contig.size = 100000; mask.regions = FALSE; and max.cluster.length.mbp = 300. SaaRclust automatically reports contigs that likely contain a misassembly and marks them as either misorientation (change in directionality of a piece of contig) or chimerism (regions of a contig that originate from different chromosomes). To reduce false positive calls, we report only misoriented and chimeric regions that are at least 400 kb and 1 Mb in length, respectively. Current version of the R package SaaRclust is available at https://github.com/daewoooo/SaaRclust (devel branch).

To evaluate large (≥ 50 kb) inversion accuracy in the final HPRC-HG002 assembly of this study we aligned Strand-seq separately to maternal and paternal haplotypes. Only chromosomes or scaffolds ≥ 1Mb were processed. We used breakpointR^72^ to detect changes in read directionality and thus putative misassemblies across all Strand-seq libraries. We concatenated all directional reads across all available Strand-seq libraries using the breakpointR function ‘synchronizeReadDir’. Next we used the breakpointR function ‘runBreakpointr’ to detect regions that are homozygous (‘ww’; ‘HOM’) or heterozygous inverted (‘wc’, ‘HET’) using following parameters: bamfile = <COMPOSITE_FILE>, pairedEndReads = FALSE, chromosomes = [chromosomes/scaffolds >= 1Mb], windowsize = 50000, binMethod = “size”, background = 0.1, minReads = 50, genoT = ’binom’. Regions designate as ‘HOM’ have the majority of reads in the minus direction, suggesting a homozygous inversion or misorientation assembly error. Those designated at ‘HET’ have roughly equal mixture of plus and minus reads, validating a true heterozygous inversion. In an ideal scenario one would expect that assembly directionality matches the directionality of Strand-seq reads and thus no homozygous inverted regions should be visible.

#### Variation benchmark analysis

We used v4.2.1 GIAB Benchmark variants with GA4GH, with v3.0 stratifications, which enabled comparative performance assessment inside and outside challenging genomic regions such as segmental duplications, homopolymers, tandem repeats^34^. Benchmarking tools from GIAB and GA4GH enable performance to be stratified by type of error (e.g., genotyping errors) and genome context (e.g., segmental duplications). Variants were first called using the dipcall assembly variant calling pipeline (https://github.com/lh3/dipcall)38. Dipcall first aligns an assembly to the GRCh38 reference genome (ftp://ftp.ncbi.nlm.nih.gov/genomes/all/GCA/000/001/405/GCA_000001405.15_GRCh38/seqs_for_align ment_pipelines.ucsc_ids/GCA_000001405.15_GRCh38_no_alt_analysis_set.fna.gz) using minimap2 (https://github.com/lh3/minimap2)73. We used optimized alignment parameters -z200000,10000 to improve alignment contiguity, as this is known to improve variant recall in regions with dense variation, such as the MHC^74^. Dipcall uses the resulting alignment to generate a bed file with haplotype coverage and call variants. All filtered variants except those with the GAP2 filter were removed. GAP2 filtered variants occurred particularly in primary-alternate assemblies in homozygous regions where the alternate contig was missing. These GAP2 variants were kept as filtered to give separate performance metrics, and treated as a homozygous variant with respect to GRCh38 by changing the genotype (GT) field from 1|. to 1|1. The resulting variant calls were benchmarked using hap.py v0.3.12 with the RTG Tools (v3.10.1) vcfeval comparison engine (https://github.com/Illumina/hap.py)39. Earlier versions of hap.py and vcfeval do not output lenient regional variant matches to the FP.al field. The hap.py comparison was performed with the v4.2.1 Genome In A Bottle HG002 small variant benchmark vcf and bed (https://ftp-trace.ncbi.nlm.nih.gov/ReferenceSamples/giab/release/AshkenazimTrio/HG002_NA24385_son/NISTv4.2.1/GRCh38/)^24^ and V3.0 of the GIAB genome stratifications (doi:10.18434/M32190). To improve reproducibility and transparency, Snakemake (https://snakemake.readthedocs.io/en/stable/)75 was used for pipeline construction and execution (https://github.com/jmcdani/HPRC-assembly-benchmarking). The extensive performance metrics output by hap.py in the extended.csv files were summarized in the following metrics for Completeness, Correctness, and Hard Regions. Completeness metrics were calculated from SNV FN rate or Recall to assess how much of the benchmark does the callset cover, where 100% means capturing all variants and 0% means capturing none. These completeness metrics were calculated at different stringencies with SNP.Recall or as a true positive (TP). ‘SNP.Recall_ignoreGT’ is a measure of how well the assembly captures at least one of the variant alleles, and is considered TP if at least one allele in a variant was called correctly, regardless of whether genotype was correct. This is calculated from ‘(SNP.TRUTH.TP + SNP.FP.gt) / SNP.TRUTH.TOTAL’ for the row with “ALL” in the FILTER column. ‘SNP.Recall’ is a measure of how well the assembly represents genotypes, and counted as TP if the variant and genotype are called correctly. When only one contig is present, we assume the region is homozygous. This is calculated from METRIC.Recall for TYPE=SNP, SUBTYPE=*, SUBSET=*, FILTER=ALL. ‘SNP.Recall.fullydiploid’is a measure of how well the assembly represents both haplotypes correctly, requiring that exactly one contig from each haplotype align to the location (contigs smaller than 10kb are ignored by dipcall by default). This is calculated from METRIC.Recall for TYPE=SNP, SUBTYPE=*, SUBSET=*, FILTER=PASS.

Correctness metrics were calculated from the false positive rate for SNVs and INDELs, converted into a phred scaled per base error rate. Each SNP and INDEL was counted as a single error on one haplotype regardless of size and genotype. ‘QV_dip_snp_indel’ is the error rate in all benchmark regions, calculated as ‘-10*log10((SNP.QUERY.FP + INDEL.QUERY.FP)/(Subset.IS_CONF.Size*2))’.

‘NoSegDup.QV_dip_snp_indel’ is the same as QV_dip_snp_indel except that it excludes segmental duplication regions.

Hard Regions metrics were calculated for particularly difficult-to-assemble regions like segmental duplications. ‘Segdup.QV_dip_snp_indel’ is the same as the ‘QV_dip_snp_indel’ Correctness metric, but only for segmental duplication regions.

To benchmark SVs, we aligned the final HG002 assembly to GRCh37 and used truvari v3.1.0 to benchmark variants against the GIAB Tier 1 v0.6 benchmark vcf in v0.6.2 benchmark regions.

#### BUSCO analyses

Busco completeness for the 41 assemblies was calculated with BUSCO v3.1.0 using the mammalia_odb9 lineage set (https://busco-archive.ezlab.org/v3/)76.

#### Annotations

Human RefSeq transcripts of type “known” (with NM or NR prefixes^77^) were queried from NCBI Entrez on December 8, 2021, and aligned to the 43 assembled haplotypes and to GRCh38 (GCF_000001405.26) and CHM13 T2T v1.1 (GCA_009914755.3). The coding transcripts and non-coding transcripts longer than 300 bp were first aligned with BLAST (evalue of 0.0001, word size 28 and best-hits options best_hit_overhang=0.1 and best_hit_score_edge=0.1) to the genomes masked with RepeatMasker (www.repeatmasker.org^78^ or Windowmasker^79^. Sets of results obtained with both masking methods were passed to the global alignment algorithm Splign^80^ (75% min exon identity, 50% min compartment identity and 20% min singleton identity) to refine the splice junctions and align exons missed by BLAST. Sequences for which no alignment with coverage higher than 95% of the query and sequences with unaligned overhangs at the 5’ or 3’ end were re-aligned with Blast and Splign to the unmasked genome, and then submitted to the same filter. Non-coding transcripts shorter than 300 bp were aligned with BLAST to the unmasked genome (evalue of 0.0001, word size 16, 98% identity and best-hits options best_hit_overhang=0.1 and best_hit_score_edge=0.1) and then with Splign (75% min exon identity, min compartment identity and min singleton identity) and submitted to the same filter as the other transcripts. The alignments for each transcript were then ranked based on identity and coverage. Transcripts that aligned best to GRCh38 sex chromosomes were filtered out prior to computing the statistics below.

Several measures for assembly completeness and correctness, and for sequence accuracy were compared across all assembly haplotypes: 1) **Unaligned genes**, either due to one or more transcript alignments being absent or too low in sequence identity; 2) **Split genes,** across two or more scaffolds; 3) **Low coverage genes,** with less than 95% of the coding sequence in the assembly; 4) **Dropped genes**, most often due to collapsed regions in the assembly; and 5) **Genes with frameshifted CDSs**, where the best-ranking alignment requires insertions or deletions to compensate for suspected insertions or deletions in the genomic sequence that cause frameshift errors. For category 4, since each RefSeq transcript is associated with a single gene^81^ and genes are not expected to overlap, unless explicitly known to, collapsed regions were identified as loci where transcripts from multiple genes co-aligned, and measured as the count of genes for which the best alignment of a transcript needed to be dropped to resolve the conflict.

### Pangenomic assessment of the assemblies

We performed pairwise alignments for all chromosomes of all 45 assemblies with the wfmash sequence aligner (https://github.com/ekg/wfmash, commit 09e73eb), requiring homologous regions at least 300 kb long and nucleotide identity of at least 98%. We used the alignment between all assemblies to build a pangenome graph with the seqwish variation graph inducer^82^ (commit ccfefb0), ignoring alignment matches shorter than 79 bp (to remove possible spurious relationships caused by short repeated homologies). To obtain chromosome-specific pangenome graphs, contigs were partitioned by aligning all of them with wfmash against the GRCh38 and CHM13 reference sequences. Graph statistics, visualizations, and pairwise Jaccard similarities and Euclidean distances between haplotypes were obtained with the ODGI toolkit^83^ (commit 67a7e5b). We performed the multidimensional analyses in the R development environment (version 3.6.3), equipped with the following packages: tidyverse (version 1.3.0), RColorBrewer (version 1.1.2), ggplot2 (version 3.3.3), ggrepel (version 0.9.1), and stats (version 3.6.3). Specifically, we applied the Classical Multidimensional Scaling on the Euclidean distance matrix to perform the Principal Component Analysis (PCA). Pangenome graphs at selected *loci* were built and visualized by using the PGGB pipeline (commit 5d26011) and the ODGI toolkit. Code and links to data resources used to build the pangenome graphs, perform the multidimensional analyses, and produce all the figures can be found at the following repository: https://github.com/AndreaGuarracino/HG002_assemblies_assessment.

### Heterozygosity analysis

To call the full spectrum of heterozygosity between the two haploid sequences, we directly compared two haploid assemblies using Mummer (v4.0.0rc1) with the parameters of “nucmer -maxmatch -l 100 -c 500”. SNP and small indels were generated by “delta-filter -m -i 90 -l 100” and followed by “dnadiff”. Several custom scripts were used to analyze the Mummer output, as described in our prior marmoset study (https://github.com/comery/marmoset). We employed SyRi (v1.5, [https://doi.org/10.1186/s13059-019-1911-0]) to detect SVs from Mummer alignments using default parameters. SVs in which more than half the feature consisted of gaps were excluded. For CNVs, we only included local tandem contractions/expansions; whole-genome copy number changes were not included in these results. To avoid false positives caused by assembly issues and insufficient detection power, we only included translocations where haplotypes reciprocally share the best alignment.

### Gene duplication analysis

Gene duplications were measured using multi-mapped gene bodies and read depth. Gencode v29 transcripts were aligned using minimap2 (version 2.17-r941) to annotate gene models. The genomic sequence of each gene was re-mapped to both HPRC-HG002 v1.0 maternal and paternal assemblies allowing for multi-mapped alignments. Gene duplications were counted as genome sequences aligned with at least 90% identity and 90% of the length of the original gene. Spurious duplications were annotated by mapping all reads back to each haplotype assembly, and filtering on low (<0.05) read depth. Code to annotate duplicated genes is available at: https://github.com/ChaissonLab/SegDupAnnotation/releases/tag/vHPRC.

### Data and Software Availability

All raw sequence data used in this study is available at the following HPRC Github: https://github.com/human-pangenomics/HG002_Data_Freeze_v1.0. The final HPRC-HG002 curated assemblies are available in NCBI under the BioProject IDs PRJNA794175 and PRJNA794172, with accession numbers GCA_021951015.1 and GCA_021950905.1, for the maternal and paternal haplotypes respectively.

## Consent

Informed consent was obtained by the Personal Genome Project, which permits open sharing of genomic data, phenotype information, and redistribution of cell lines and derived products.

## Author Contributions

E.D.J. Co-coordinated evaluation, performed analyses, and wrote paper

K.H. Co-coordinated evaluation and curation

K.H.M. Overall coordination A.Rh. Mercury metric evaluations

T.L. Assembly validations

V.A.S. Genome alignment, annotation/frameshift issues

B.P. ONT assemblies and overall coordination

M.H. Shasta assembly, CAT-annotation, QUAST

J.M.Z. Variant analyses

I.M.H. Variant analyses A.Re. Variant analyses

E.E.E. SDA, collapsed duplications, Strand-seq analyses

M.R.V. SDA, collapsed duplications, Strand-seq analyses

D.Po. asm14, assembly inversion validation, Strand-seq analyses

W.H. Bioinformatic support

H.Li Genome assembly and overall analyses

S.G. Genome assembly and phasing evaluation E.Ga. Pangenome alignments, Jaccard, PCA, and MHC

M.E. PacBio HiFi adaptor assessments

F.T-N Annotations, gene loss, and associated analyses J.Wo. Curation

J.C. Curation

J.T. Curation

J.M. Variant analyses

N.D.O. Variant analyses J.Wag. Variant analyses

M.A. Genome assemblies and titration

A.G. Pangenome alignments, Jaccard, PCA, and MHC

G.Z. Heterozygosity circos plot analyses supervision

C.Y. Heterozygosity circos plot analyses

A.T. Curation of final diploid HG002 assemblies

G.A.L. Centromere and telomere analyses

G.F. asm23 and final HRPC HG002 references

H.C. asm9

S.K. asm 6, 7, 19, 20, and MHC evaluation

A.M.P. Assembly coordination

J.G. asm2 and asm22

T.M. asm14

P.E. asm14

M.C.S. asm17 M.Ki. asm17

C-S.C asm3 and asm4

J.K. asm16

A.W. asm16

S.L.S. asm1 and asm15

J.R. asm13

F.N. asm12

C.X. asm12 J.Wan. asm12

F.L. asm12

M.C. HG002 reference segmental duplication expansions

P.C. Shasta assembly

T.P. Shasta assembly, scaffolding and polishing

K.S. Shasta assembly, scaffolding and polishing

O.F. Bionano analyses

A.V.Z. asm1 and asm15 A.Sh. asm1

D.Pu asm1

M.Ko. Flye haploid assemblies

J.Lu. Sample processing, analyses, and submissions

M.J. Nanopore data generation, analysis, data management E.Gr. Dovetail Genmonics Hi-C

S.S. Arima Genomics Hi-C A.Sc. Arima Genomics Hi-C

R.S.F. PacBio production activities, evaluation methods

H.E.O. Nanopore data generation

A.D.S Strand-seq data production

J.O.K. Strand-seq data production

P.H. Strand-seq data production

C.S. Strand-seq data production

A.H. Bionano diploid assembly maps J.Le. Bionano diploid assembly maps

D.H. HPRC coordination

T.W. HPRC coordination

L.L.F. Logistics, data release, project support

D.L. Logistics, data release, support

S.C. Logistics, data release, support

### Competing Interest

E.E.E. is a scientific advisory board (SAB) member of Variant Bio, Inc. J.L. and A.H. are former and current employees and shareholders of Bionano Genomics, Inc respectively.

## Funding

The primary source of funding for this study was from NHGRI grant U01HG010971 (https://grants.nih.gov/grants/guide/rfa-files/rfa-hg-19-004.html). Additional funding: HHMI (E.D.J.), NHGRI grants HG002385 and HG010169 (E.E.E.). The computational resources and personnel support for: the DipAsm assemblies were HG010906/HG/NHGRI NIH HHS and NNF21OC0069089 to SG; NCI U01CA253481 to MCS; and NSF DBI-1627442 to MCS. T.M. and P.E. (asm14 team) acknowledge the support by the BMBF-funded de.NBI Cloud within the German Network for Bioinformatics Infrastructure (de.NBI) (031A532B, 031A533A, 031A533B, 031A534A, 031A535A, 031A537A, 031A537B, 031A537C, 031A537D, 031A538A), and the German Research Foundation (391137747 to T.M.). E.D.J, E.E.E. D.H. are investigators of the Howard Hughes Medical Institute. This work was supported in part by the National Center for Biotechnology Information of the National Library of Medicine (NLM), National Institutes of Health, and by National Institute of Standards and Technology intramural research funding.

## Acknowledgements

Certain commercial equipment, instruments, or materials are identified to specify adequately experimental conditions or reported results. Such identification does not imply recommendation or endorsement by the National Institute of Standards and Technology, nor does it imply that the equipment, instruments, or materials identified are necessarily the best available for the purpose. T.M. and P.E. (asm14 team) acknowledge the computational infrastructure and support provided by the Centre for Information and Media Technology at Heinrich Heine University Düsseldorf. We thank Peter Audano of the Eichler lab for bioinformatic support.

## Supplementary Notes

**Supplementary Note 1**: Estimation of diversity representation

Based on theoretical estimates from SNPs in the 1000G project, and using Kruglyak et al 2000 formula^84^, to detect 100% of minor alleles greater than 2% in the human world population or ∼97% of those above 1%, one would need to generate complete sequence for ∼350 diploid genomes (700 haplotypes) from diverse populations, the number thus far funded. To obtain 99% of of those above 1%, one would need ∼450 genomes (900 haplotypes). Minor allele frequency is the frequency at which the second most common allele occurs in a population. This calculation will differ for structural variants, estimated at 50 genomes for alleles above 5%. These values will need to be revised in an iterative fashion, as more relatively complete and error-free genomes are generated. For example, as validated in this study, haplotype diversity is uneven across the genome, e.g. where centromeres have much greater diversity than most other genic parts of the genome. Also the technology used will make a difference. These values were determined based mostly on short-read only assemblies, which do not assemble centromeres well, and also not GC-rich or repeat regions well.

Supplementary Note 2: Additional reads with adaptors that needed to be removed.

After the standard method of removing adaptors in the sequence reads, we screened the presence of any remaining adaptors in the HG002 Pacbio libraries. We found that 0.3-0.4% of reads (∼2,000) still contained at least 1 adapter per SMRT cell movie, and that adds up to a 1.7% frequency overall (all contaminated reads/all reads). Such adaptor contaminated reads were greatly reduced with chemistry 3.0 relative to 2.0 and pre2.0. The PacBio 45 bp blunt adapter was the sequence that was not properly clip in the sequence processing: > gnl|uv|NGB00972.1:1-45 Pacific Biosciences Blunt Adapter ATCTCTCTCTTTTCCTCCTCCTCCGTTGTTGTTGTTGAGAGAGAT These were due to reads with adapter 2-dimers (more prevalent in chem pre2.0 than in chem2.0). The location of the adapters were mostly (90%) at the beginning of the reads (first 1/3rd of the read length), and 87% within the first 100bp; 0.1% in the middle of the read (2/3 of the read); and the remaining 9.5% at the end of the read (last third of the read), and 9.3 % within the last 100bp. To identify reads with reads of all these categories, we used HiFiAdapterFilt (https://github.com/sheinasim/HiFiAdapterFilt). Some reads were entirely made of adaptors, which presumably occurs by self ligation in the library preparation. We removed all such reads with adaptors for the final HG002 assembly, even if they had human sequence attached, because using cut versions of these reads led to lower assembly metrics relative to without them. Later we switched to CutAdapt^35^, as the results had about 90% overlap with HiFiAdapterFilt, but Cutadapt was about 2 times faster and slightly more sensitive (5-10%).

## Extended Data Figures

**Extended Data Fig. 1.**
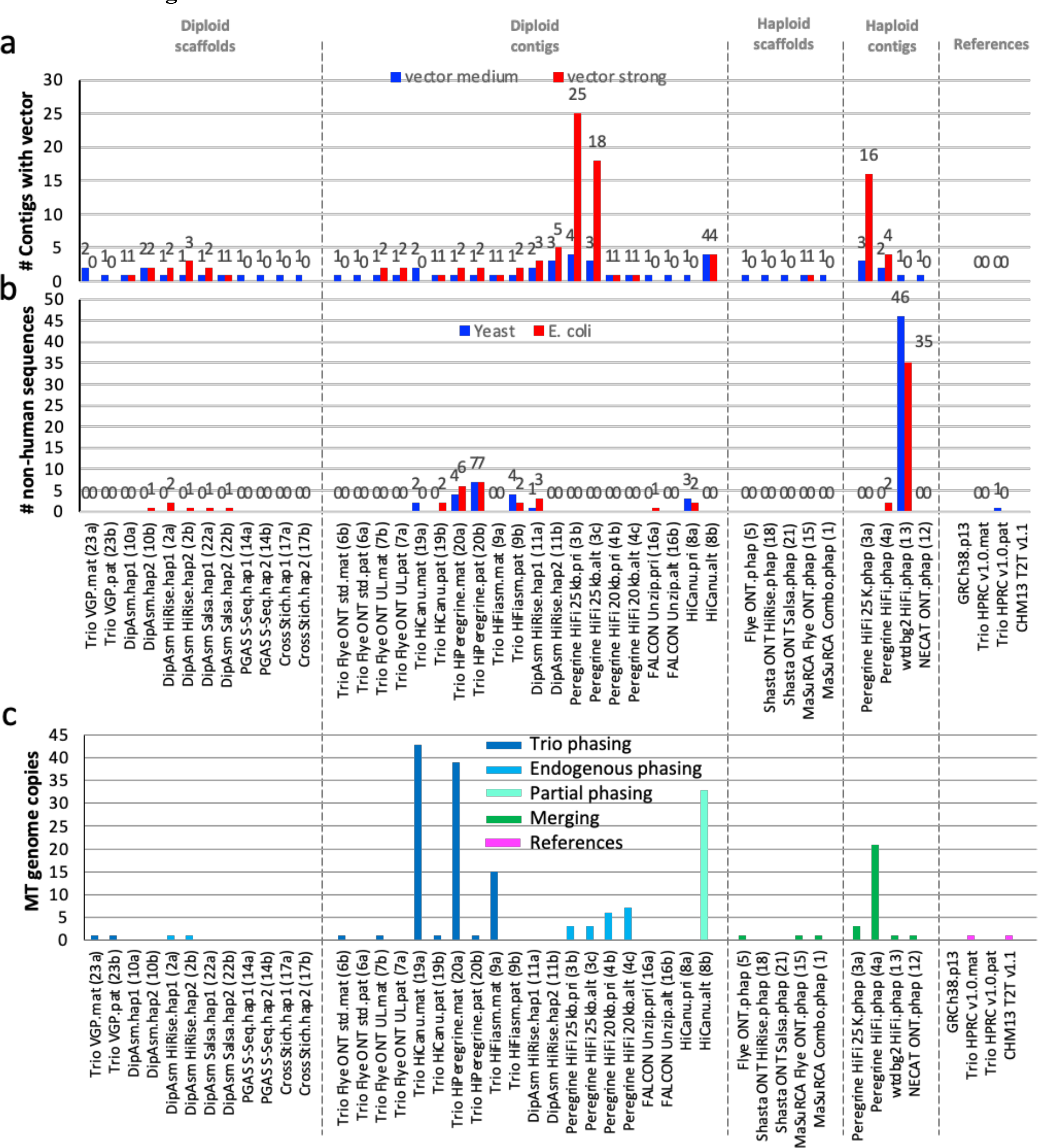
Non-human and organelle genomes found in the human genome assemblies. **a**, The number of contigs that had remaining library clone vector sequences in each assembly. Medium used a blastn score 19-29; strong a score > 30 https://www.ncbi.nlm.nih.gov/tools/vecscreen/about/. **b**,The number of contigs with non- human yeast and *E.coli* sequences. Values above columns are the specific numbers. **c**, The number of endogenous mitochondrial genome copies found in each assembly.

**Extended Data Fig. 2.**
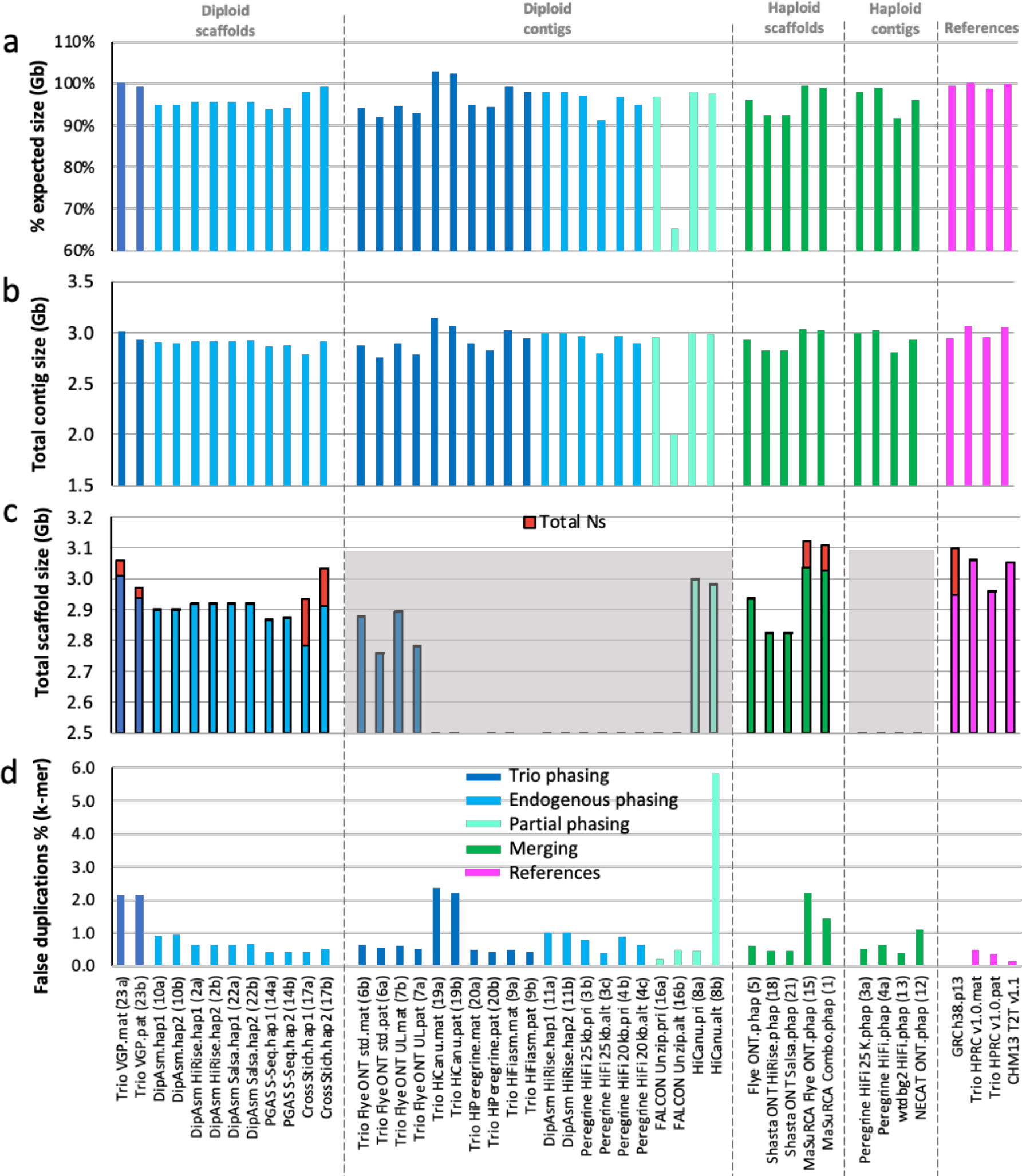
Assembly size and false duplication metrics. **a,** Percent assembly sizes of expected maternal with Chr X (3,054,832,041 bp) or paternal with Y (2,995,432,041 bp) for trio-based assemblies, or simply relative to maternal size for all other assemblies**. b,** Total summed length of all contigs**. c**, Total summed length of scaffolds, with proportion contributed by Ns (red) in gaps. **d**, estimated percent of assembly size that is due to false duplications based on k-mer values for each haplotype.

**Extended Data Fig. 3.**
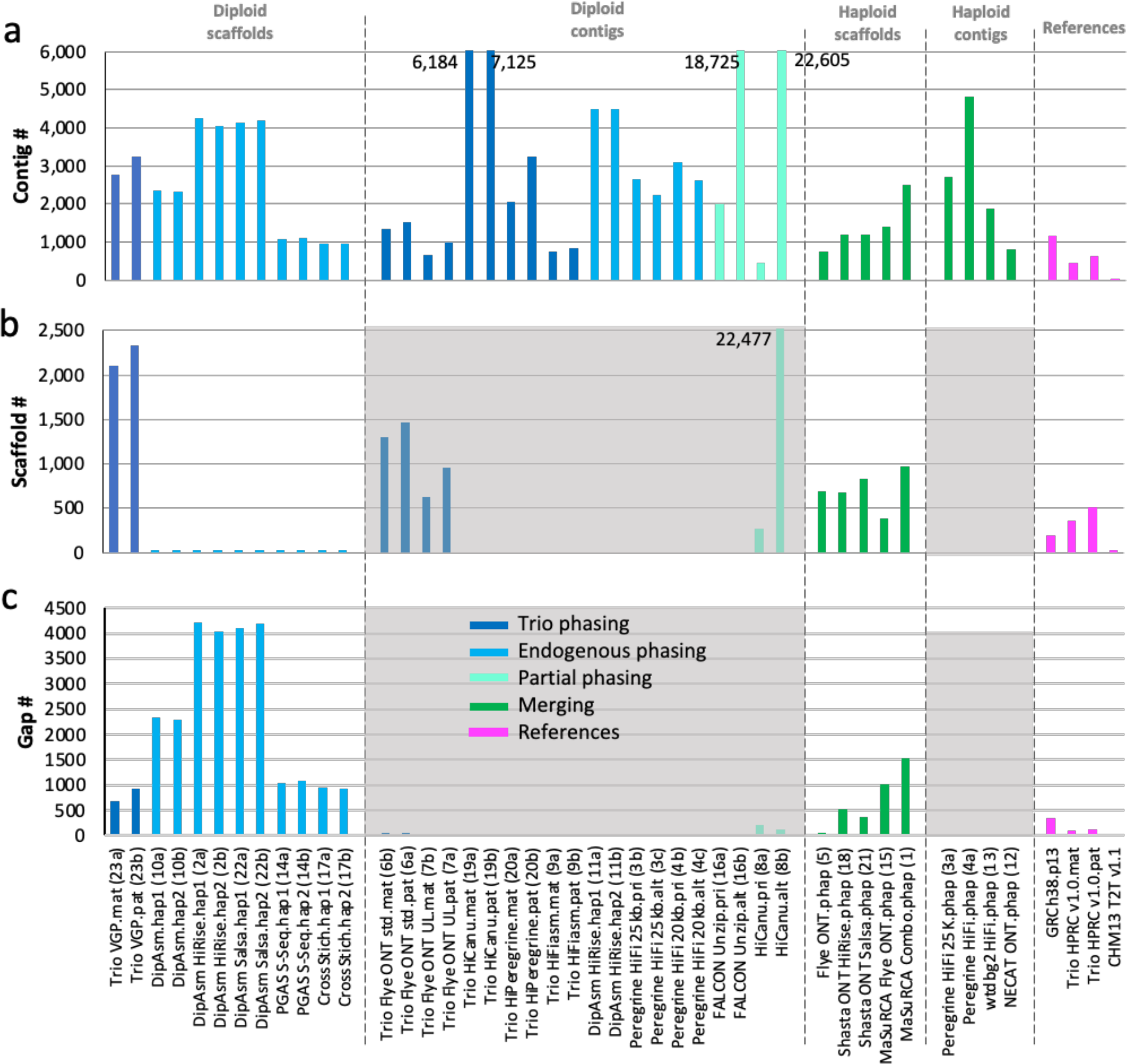
Contig, scaffold, and gap metrics. **a,** Percent assembly sizes of expected maternal with Chr X (3,054,832,041 bp) or paternal with Y (2,995,432,041 bp) for trio-based assemblies, or simply relative to maternal size for all other assemblies**. b,** Total summed length of all contigs**. c**, Total summed length of scaffolds, with proportion contributed by Ns (red) in gaps. **d**, estimated percent of assembly size that is due to false duplications based on k-mer values for each haplotype. Values above the maximum on the y-axis are written in the graph so as to not visually scale down the majority of the results.

**Extended Data Fig. 4.**
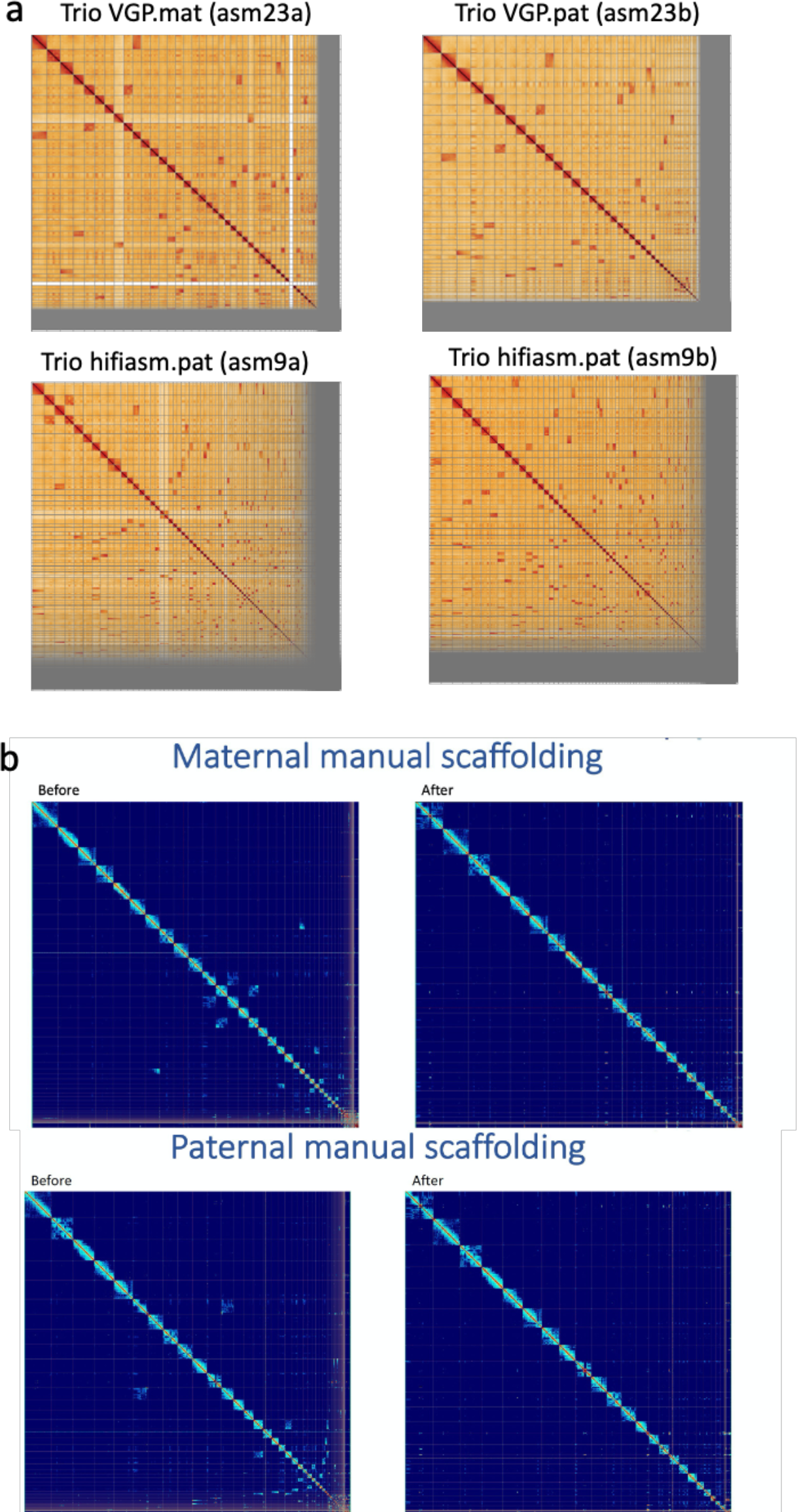
Hi-C contact maps. **a,** Example Hi-C contact maps for bakeoff maternal (mat) and paternal (pat) haplotype assemblies. The Trio VGP scaffolded assembly has several dozen large joins and many small ones to make from the off-diagonal signals. The Trio hifiasm contig only assembly as expected has many more needed. **b,** Reference HPRC HG002 assemblies for each haplotype before and after manual curation, showing less off diagonal signals and no major scaffolds/contigs not placed in chromosomes after curation.

**Extended Data Fig. 5.**
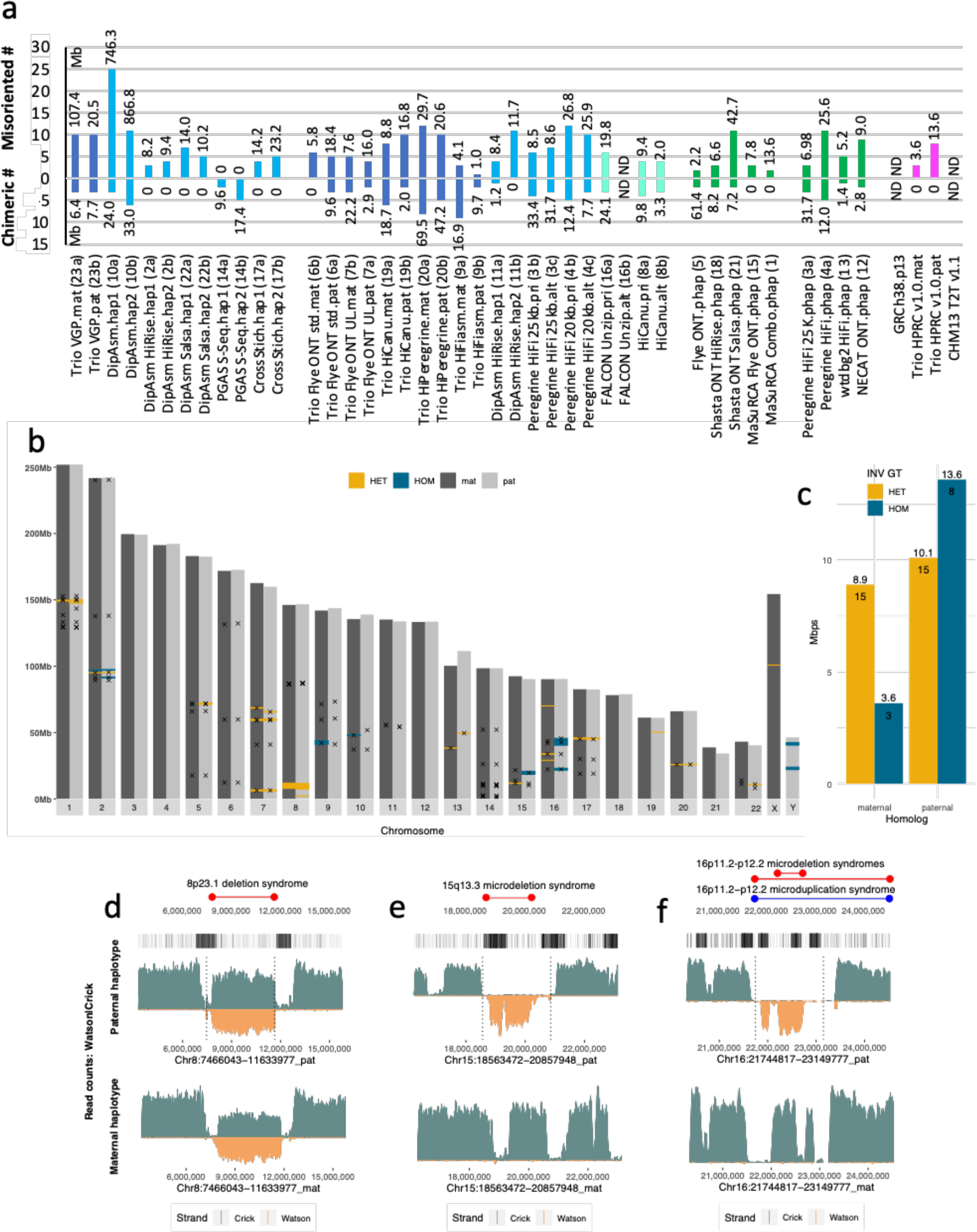
Strand-seq validations. **a**, Total number and total Mb of chimeric and misorientation errors for each assembly according to Strand-seq validations. **b**, Chromosome overview of large (> 50Kbp) Strand-seq supported and unsupported inversions. ‘x’, location of inversions (n=59) between haplotypes of the final HG002 assembly. HET, regions with roughly equal mixture of plus (Crick) and minus (Watson) Strand-seq reads supporting the heterozygous inversions (yellow, n=30). HOM, regions with Strand-seq reads mapped to the opposite orientation in disagreement with heterozygous inversions and thus a possible assembly error (HOM, blue, n=11). **c**, Barplot of total size and total number of regions genotyped as HET and HOM validated inversions. **d-f**, Example cases where the heterozygous assembly inversion matched (**d**) or did not match (**e,f**) the expected direction of the Strand-seq reads in the final HG002 assembly. Top track: Positions of known morbid CNVs (red – deletions, blue – duplications). Second track: Position of segmental duplicated sequences (black marks - DupMasker) in the paternal assembly. Third and fourth track: Coverage of Strand-seq reads aligned to the paternal and maternal assemblies of HG002 (binsize: 50kb, stepsize: 1kb) with Crick read counts (teal, above) and Watson read counts (below, orange). Regions with roughly equal coverage of Watson and Crick count represent a validated heterozygous inversion as only one homolog is inverted in respect to the de novo assembly (**d**). Regions with reads aligned only in Watson orientation represent an assembly error because assembly directionality does not match Strand-seq read directionality (**e,f**). Vertical dotted lines highlight the predicted breakpoints of assembly errors as well as predicted heterozygous inversion.

**Extended Data Fig. 6.**
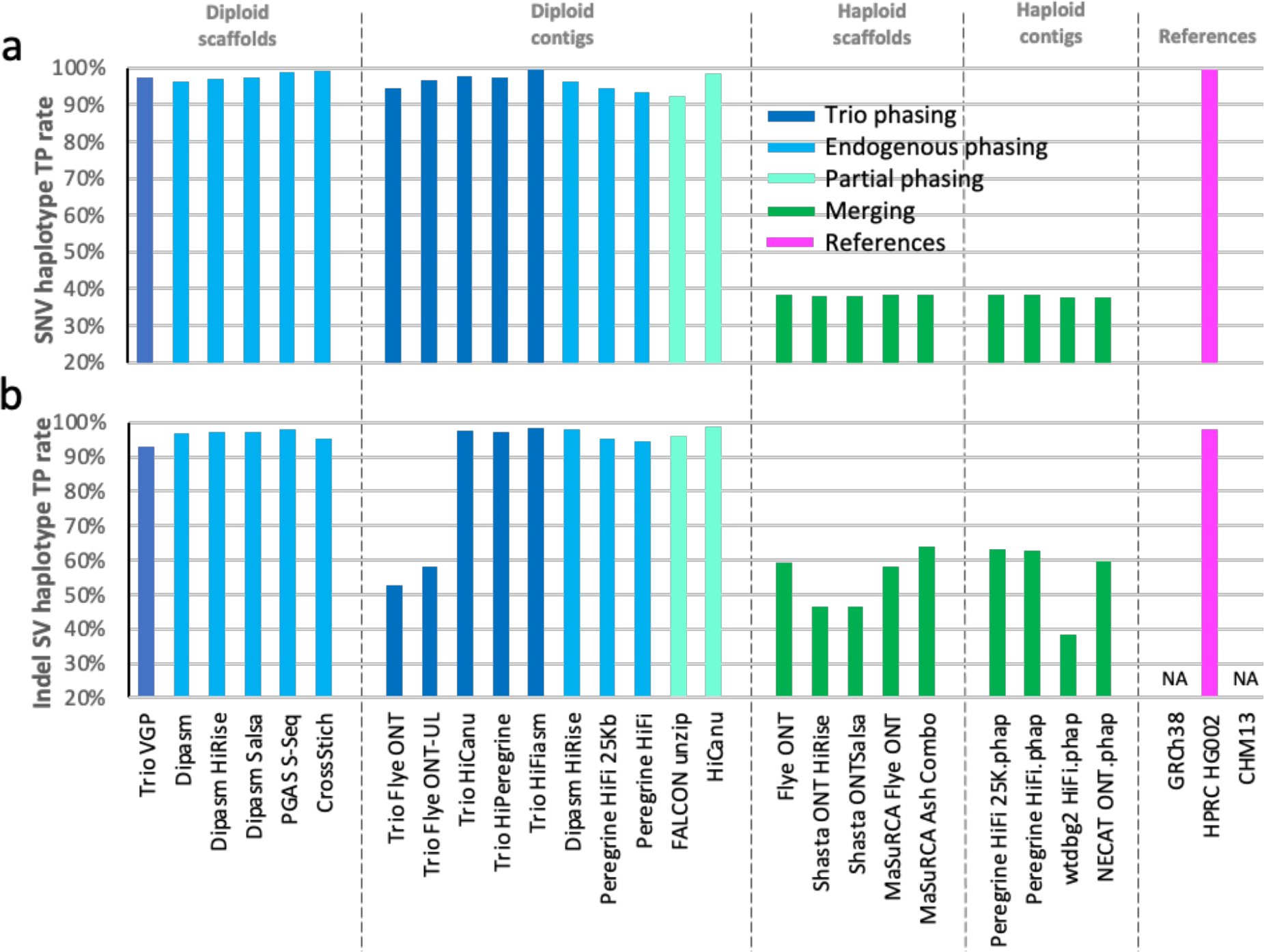
Variant benchmarking. **a**, True positive percent of known SNVs found between HG002 haplotypes in each assembly. **b**, True positive percent of known small indels found between HG002 haplotypes in each assembly. For the diploid assemblies, comparisons were made between the two haplotypes (maternal vs paternal for the trio assemblies; haplotype 1 vs haplotype 2 for the non-trio assemblies). For the haploid assemblies, we scored as TP if at least one of the variants were found.

**Extended Data Fig. 7.**
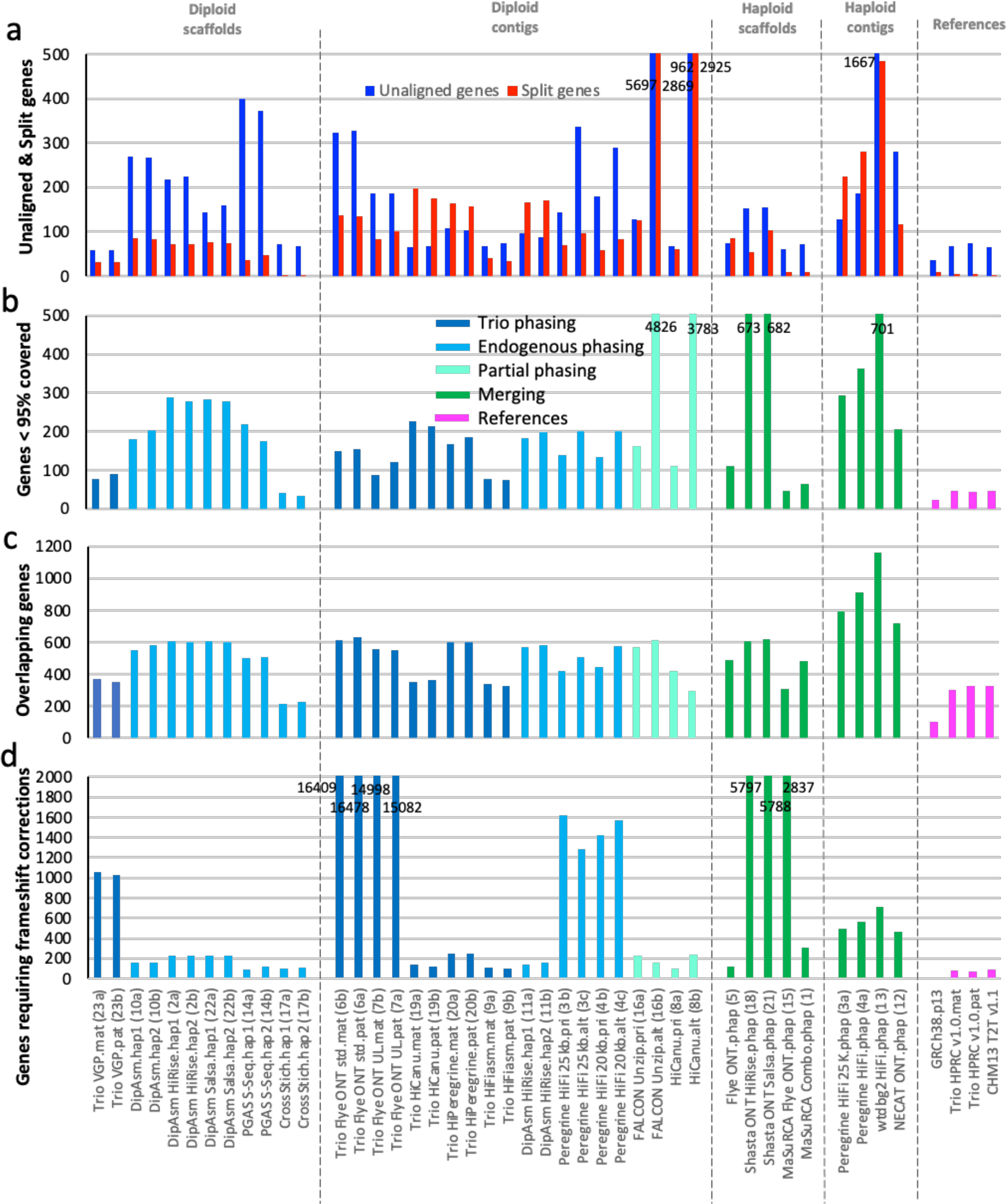
Annotation benchmarking. **a**, Side-by-side comparisons of gene transcripts that did not align to each assembly (blue) versus those that were split between two or more scaffolds/contigs (red). **b**, Number of genes that had less than 95% the length covered in the assembly. **c**, Genes in the assemblies with overlapping transcripts due to possible collapse in the assemblies. **d**, Genes requiring frameshift corrections to make a complete protein. Values written in the graphs for those off the chart, in order to not mask the lower values of most other assemblies.

**Extended Data Fig. 8.**
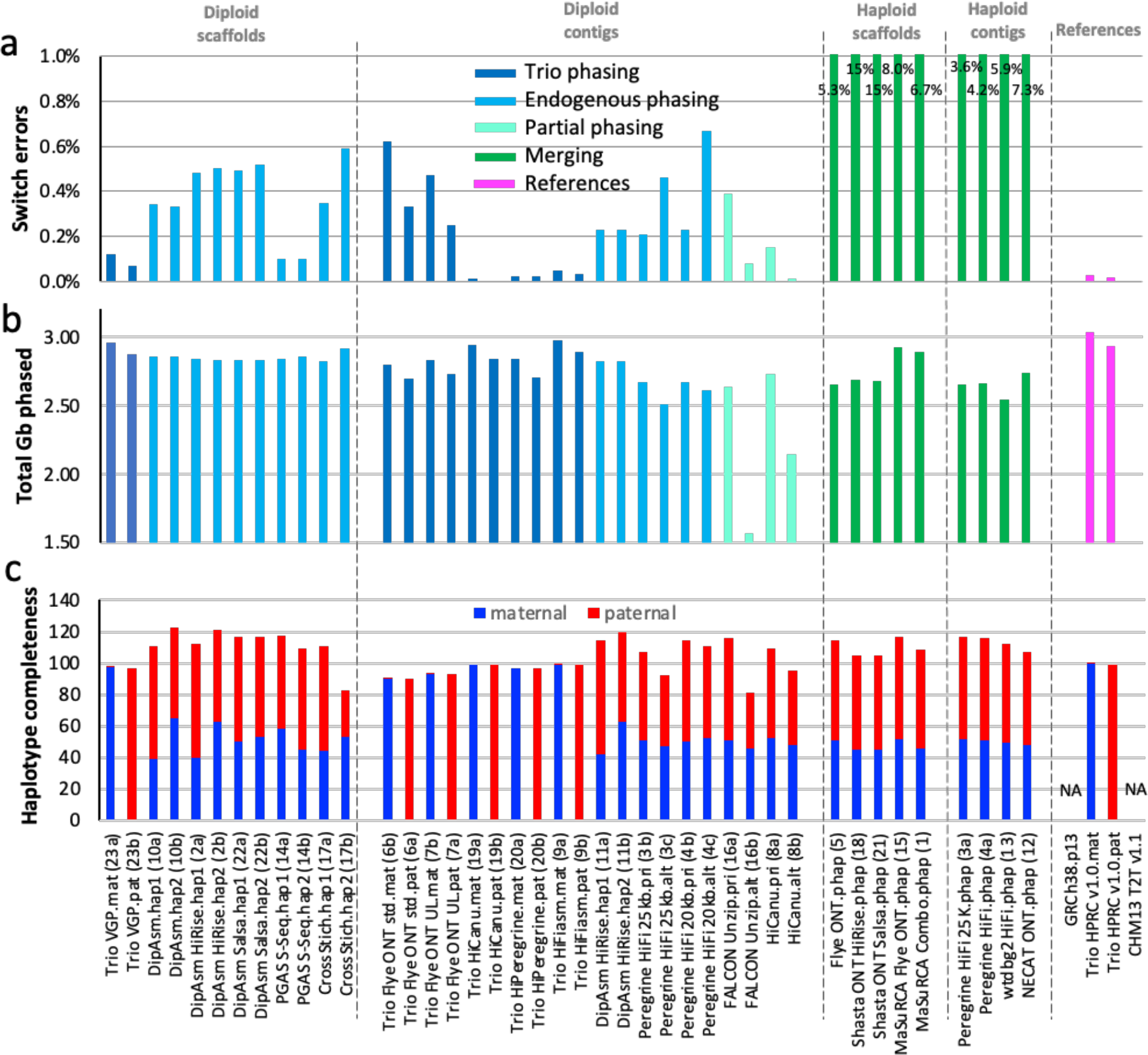
Haplotype phasing metrics. **a**, Haplotype switch errors within scaffolds and/or contigs of each assembly (lower % is more accurate). Values written in the graphs for the haploid assemblies (greens) are off the chart, in order to not mask the lower switch error values of most other assemblies. **b**, Total Gb of each assembly that has been haplotype phased (∼3.0 is the theoretical maximum of the maternal haplotype; 2.9 for the paternal). **c**, Haplotype phasing completeness according to parental k-mer statistics for each assembly. A complete phased assembly will have both maternal (blue) and paternal (red) each at 100% without mixture from the other. The trio approaches had nearly full phase separation, whereas the non-trio approaches nearly had half and half separation because there was not an attempt to phase across contigs or scaffolds/chromosomes belonging to the same maternal or paternal haplotypes. Combined values over 100% indicate a mixture of haplotype presumably due to false duplications; although values under 100% could still have false duplications.

**Extended Data Fig. 9.**
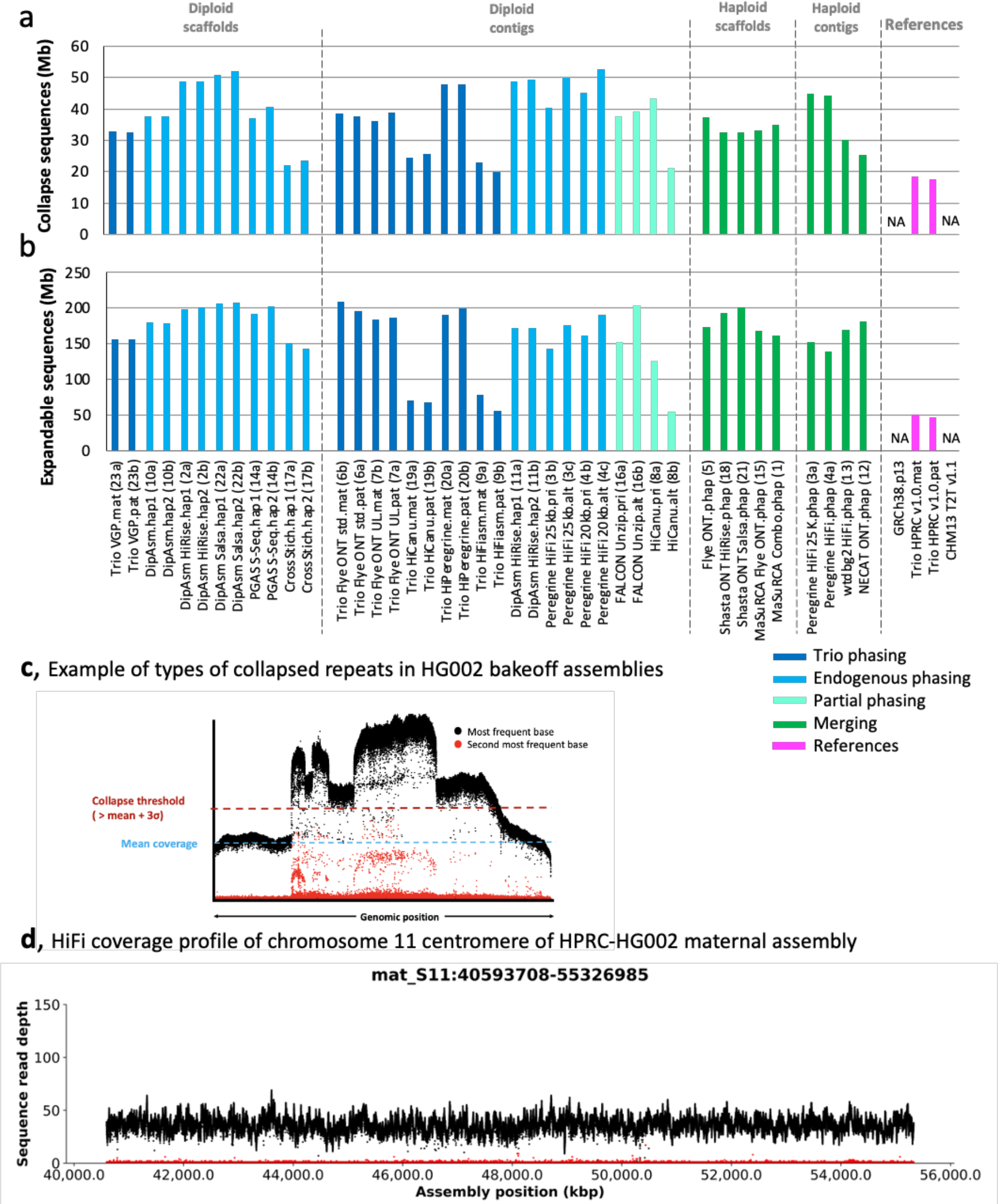
Collapsed sequence metrics. **a**, Estimated amount of bp that are collapsed in each assembly (smaller is better). Collapses are most often due to repetitive sequences. **b**, Estimated amount of bp that are potentially expandable. The smaller, the more accurate the assembly. We estimate that most of these collapses are in centromeric regions and satellites, with a smaller proportion coming from segmental duplications. Abbreviations and color coding explanations are the same as in Fig. 1 legend. **c**, Example collapse region of one of the HG002 assemblies, where read coverage pile up in the collapsed region is two or more times higher than the mean coverage of the genome. **d**, Example of HiFi read coverage across and assembled centromere, of HG002 maternal Chr 11, showing no evidence of collapsed repeats or coverage dropouts.

**Extended Data Fig. 10.**
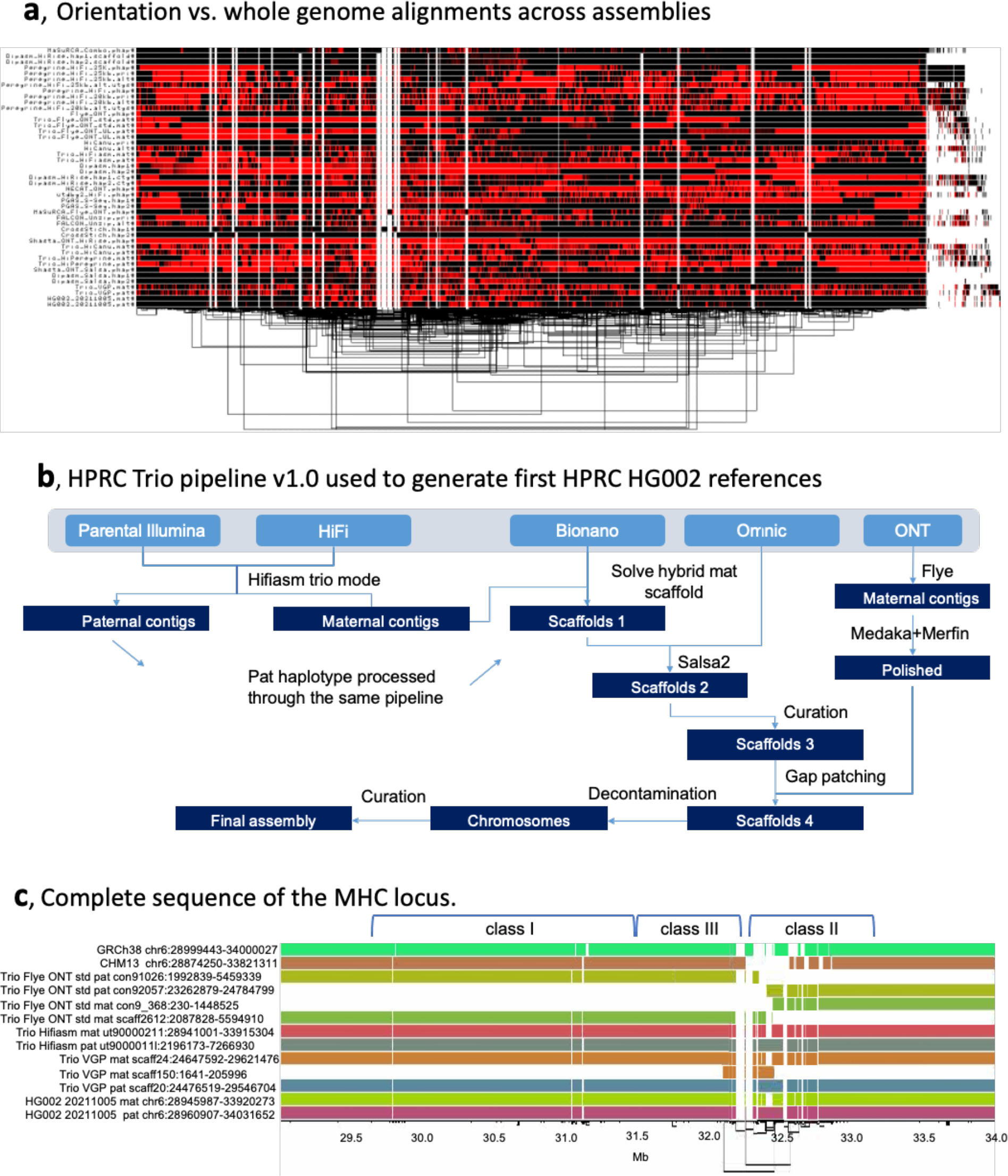
Pangenome alignment and generation of high-quality HPRC-HG002 v1.0 dioploid assemblies. **a**, Output of graph-based alignment of all chromosomes concatenated from all 45 HG002 assemblies of this study (both haplotypes of diploid assemblies). Red vs Black, are same orientation. Connections at the bottom, is a clustering of the alignments. **b**, HPRC v1.0 pipeline developed to produce the reference quality HPRC-HG002 v1.0 maternal and paternal assemblies of this study. All steps shown are highlighted for the maternal data, but the paternal data went through the same pipeline. **c**, Graph-based alignment of a 5 Mb region of human chr 6 containing the MHC locus of the Trio-based assemblies and GRCh38 and CHM13 references. Each color is a different assembled haplotype. Note the Trio hifiasm assembly and the final HG002 assembly that used Trio hifiasm assembled the entire MHC locus into one single contig.

**Extended Data Fig. 11.**
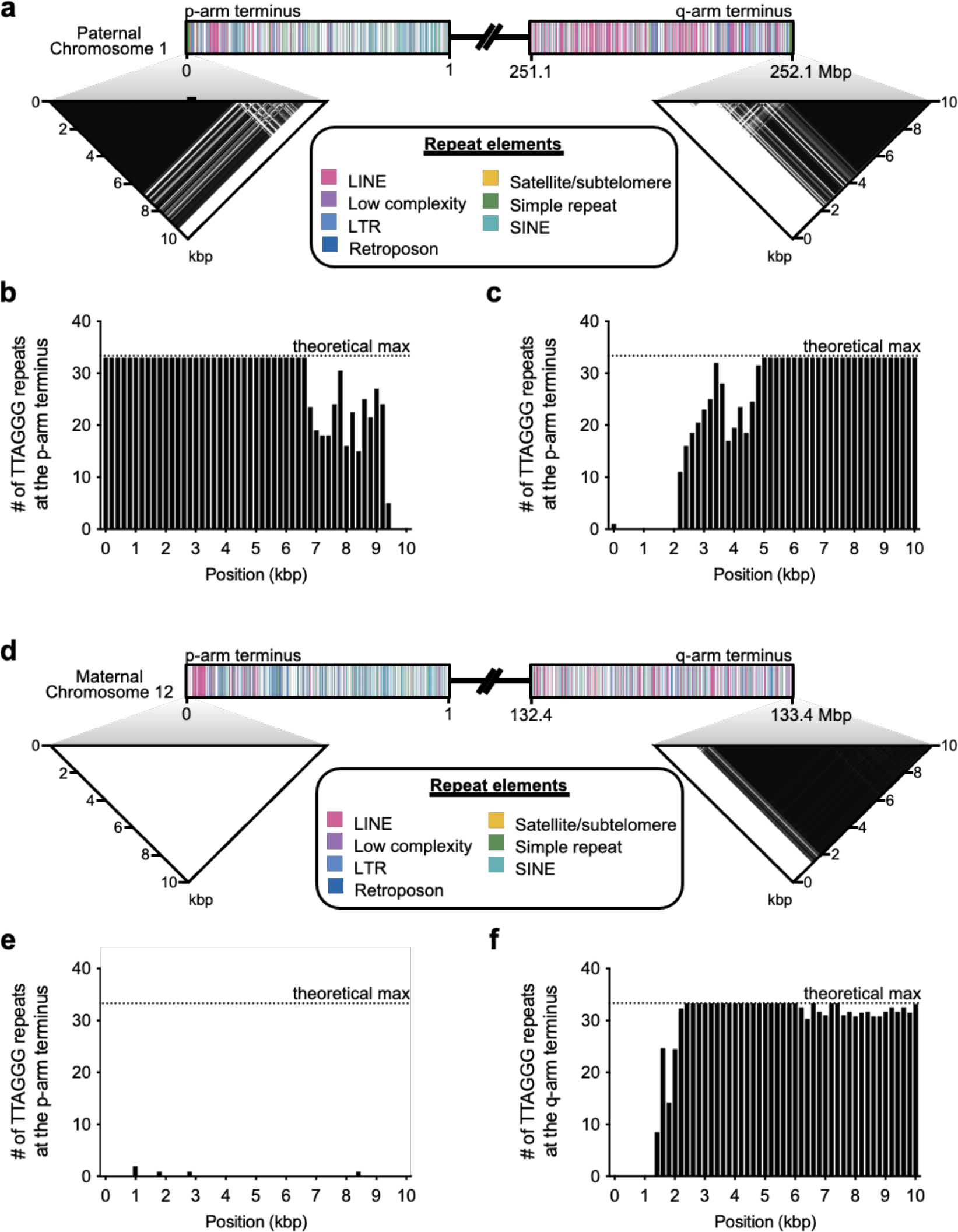
Example presence of telomeres. **a,** Telomere repeats within 10 kbp of each arm of HG002 Chr 1, paternal haplotype. The darker the density the higher the repeat copy number. **b-c**, Density of telomere repeats for each arm, in 200 bp bins. 33 x 6bp repeats is the theoretical maximum per 200 bp. d, Telomere repeats within 10 kbp only found for the q-arm of HG002 Chr 12, paternal haplotype. **e-f**, Canonical pattern of the telomere repeats only found in the q-arm of the assembly. Color coding, the different types of repeats found within 1 Mb of each arm. The similar patterns between Chr 1 and 12 indicate that only the q-arm telomere is missing from Chr 12.

## Supplementary Figures

**Supplementary Fig. 1.**
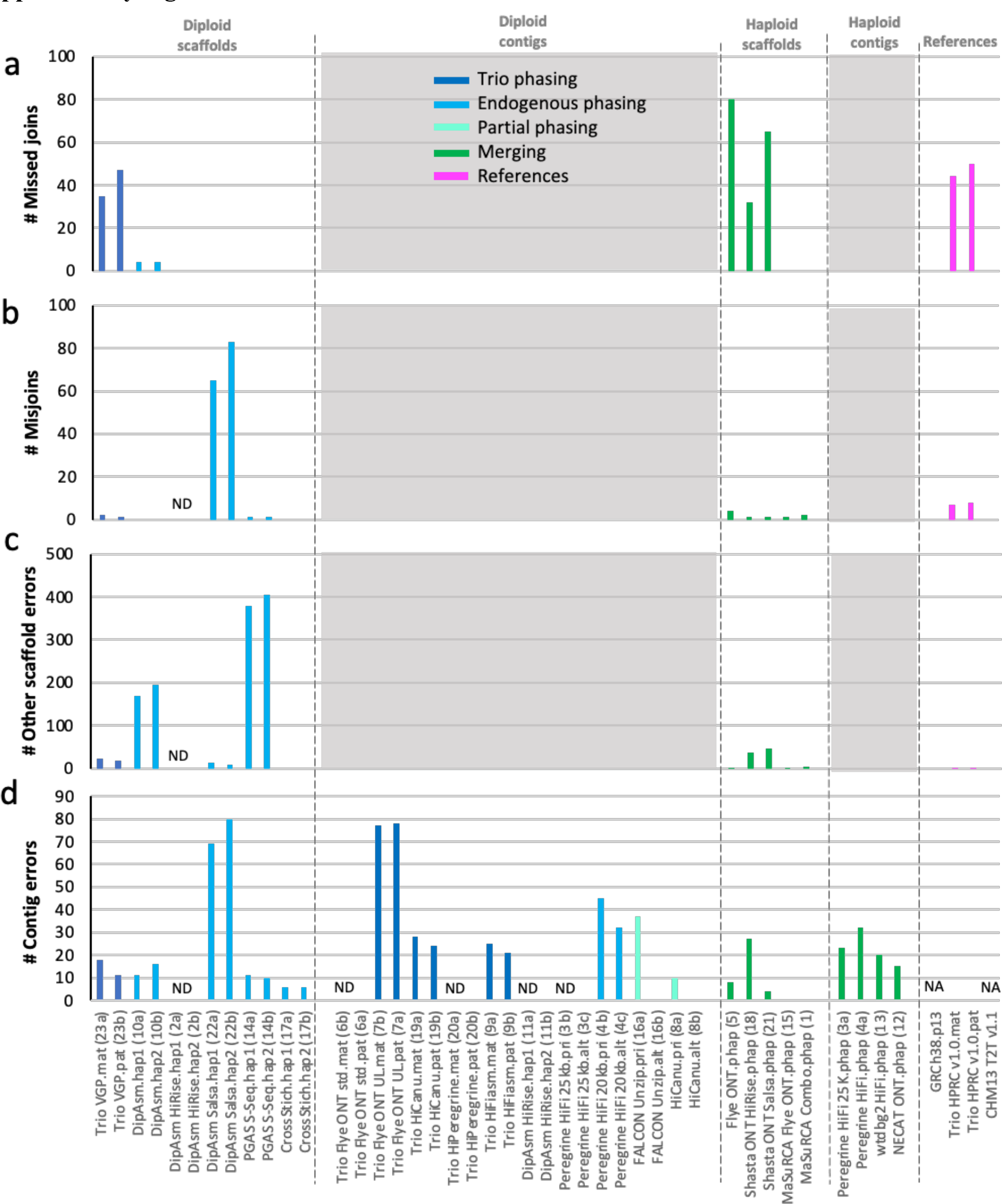
Curation structural analyses. **a**, Missed contig joins that should have been brought together in the same scaffold. **b**, Misjoins of contigs that should not have been brought together. **c**, other scaffold errors, including false inversions and translocations. **d**, Contig errors, including chimeric contigs. ND = not done, as it would be redundant with another assembly approach. NA = Not applicable, as these references were curated by a different process, with nevertheless many errors corrected.

**Supplementary Fig. 2.**
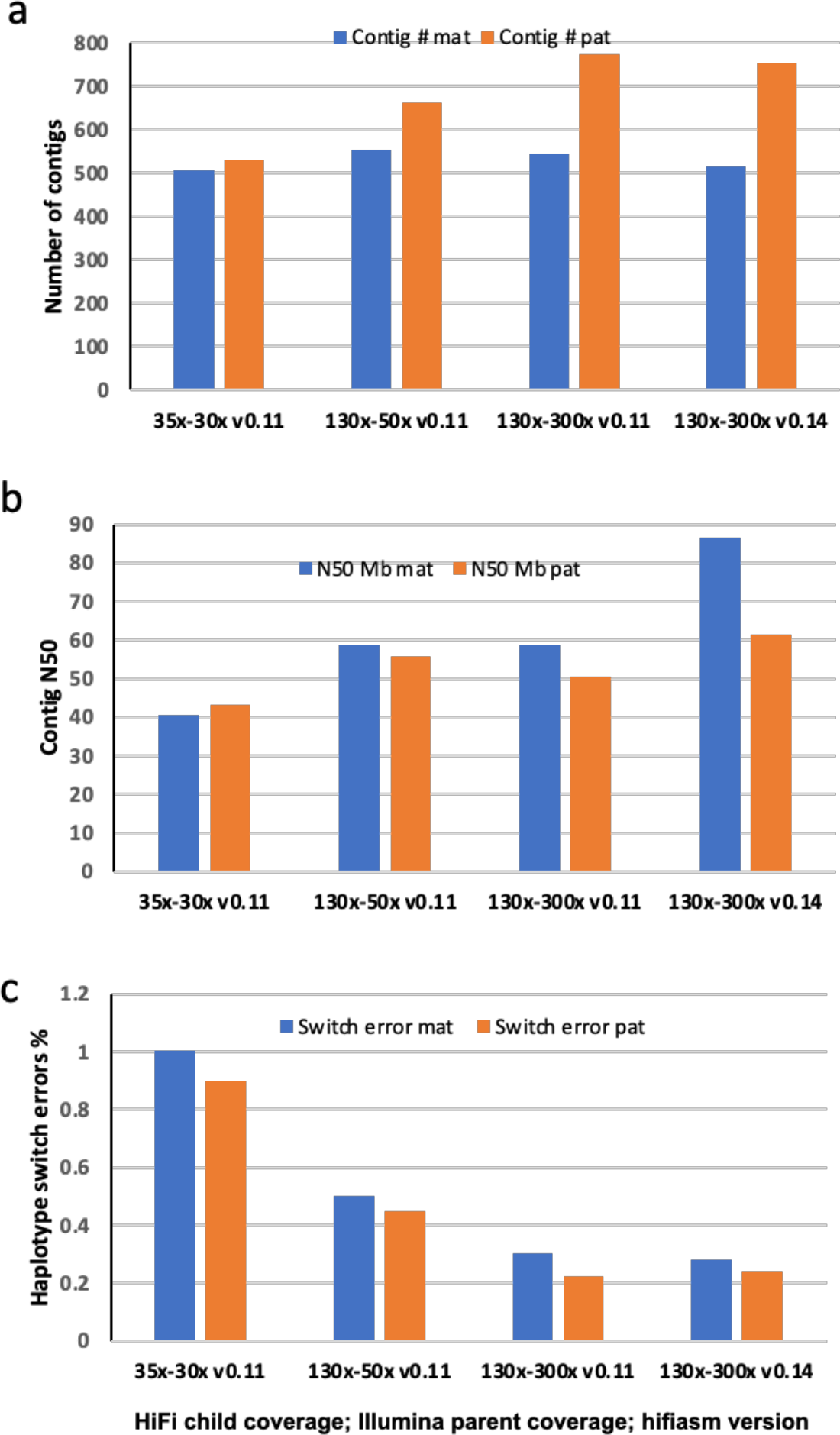
Titration of read coverage and hifiasm version on three assembly metrics. **a**, Number of contigs for each haplotype. **b**, Contig N50 for each haplotype. **c**, Haplotype switch errors for each haplotype. Two HiFi coverage levels were tested: 35x and the near full 130x data set with different library insert sizes. Three parental Illumina coverage levels were tested: 30x; 50x; and 300x. Two trio hifiasm versions were tested: v0.11 and 0.14. The increase in the number of contigs with increasing child and parental coverage could be due to more complete contigs with diverse centromere sequences, human EBV virus, and mitochondrial genomes. Hifiasm v0.14 increased contig N50. Increased child or parental sequence coverage decreased haplotype switch errors. The 130x-300x v0.14 assembly was used for the final HPRC-HG002 assembly of this study.

**Supplementary Fig. 3.**
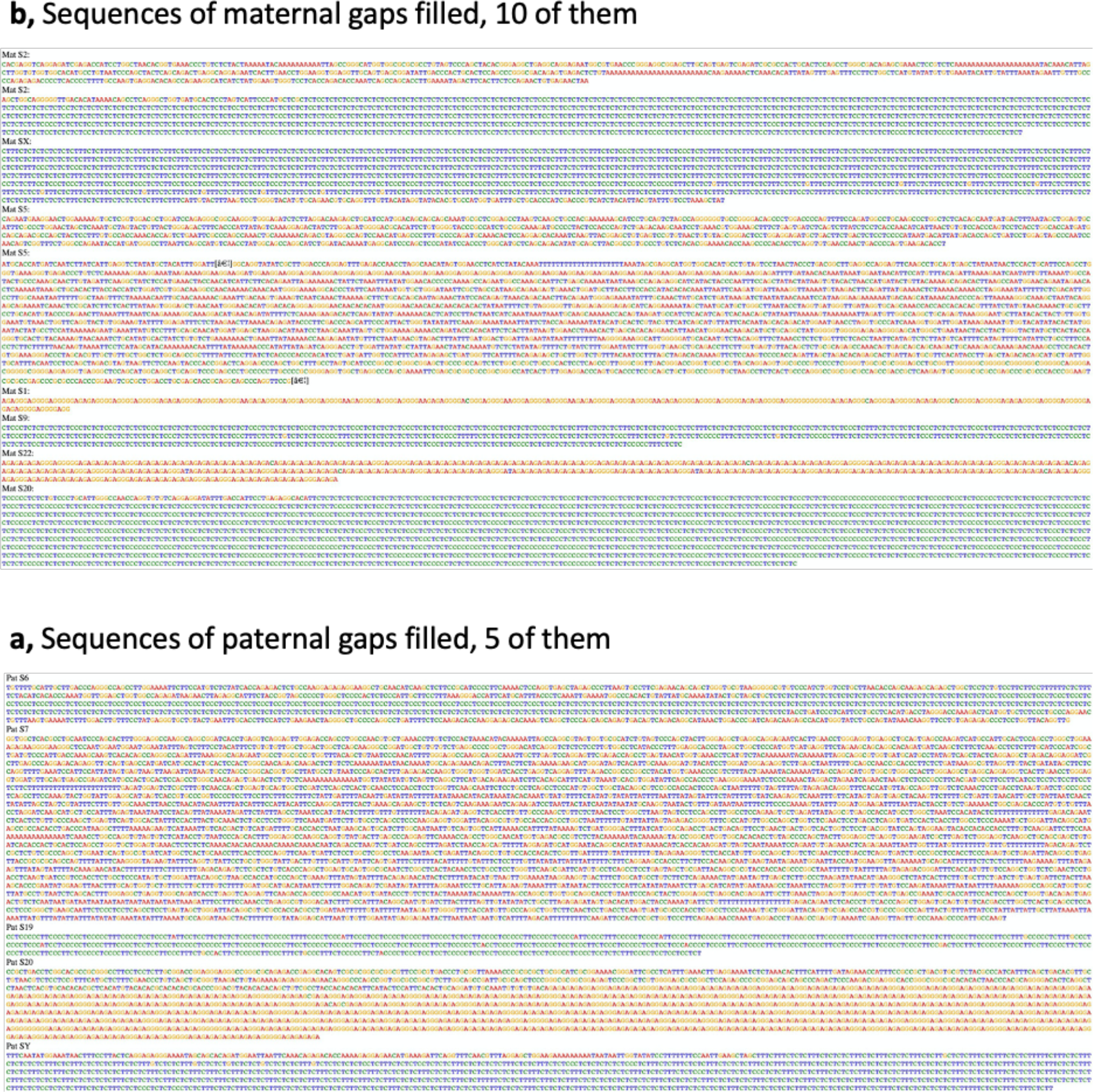
Gap filled sequences. **a**, Sequences filled in the maternal scaffolds. **b**, Sequences filled in the paternal scaffolds. Only the sequences used in the gap space are shown. These gaps between Pacbio HiFi-based contigs in scaffolds were filled with ONT-based contigs. Orange = G; Red = A; Green = C; Blue = T. Notice the GA or TC repeats in most of the filled gap sequence.

**Supplementary Fig. 4.**
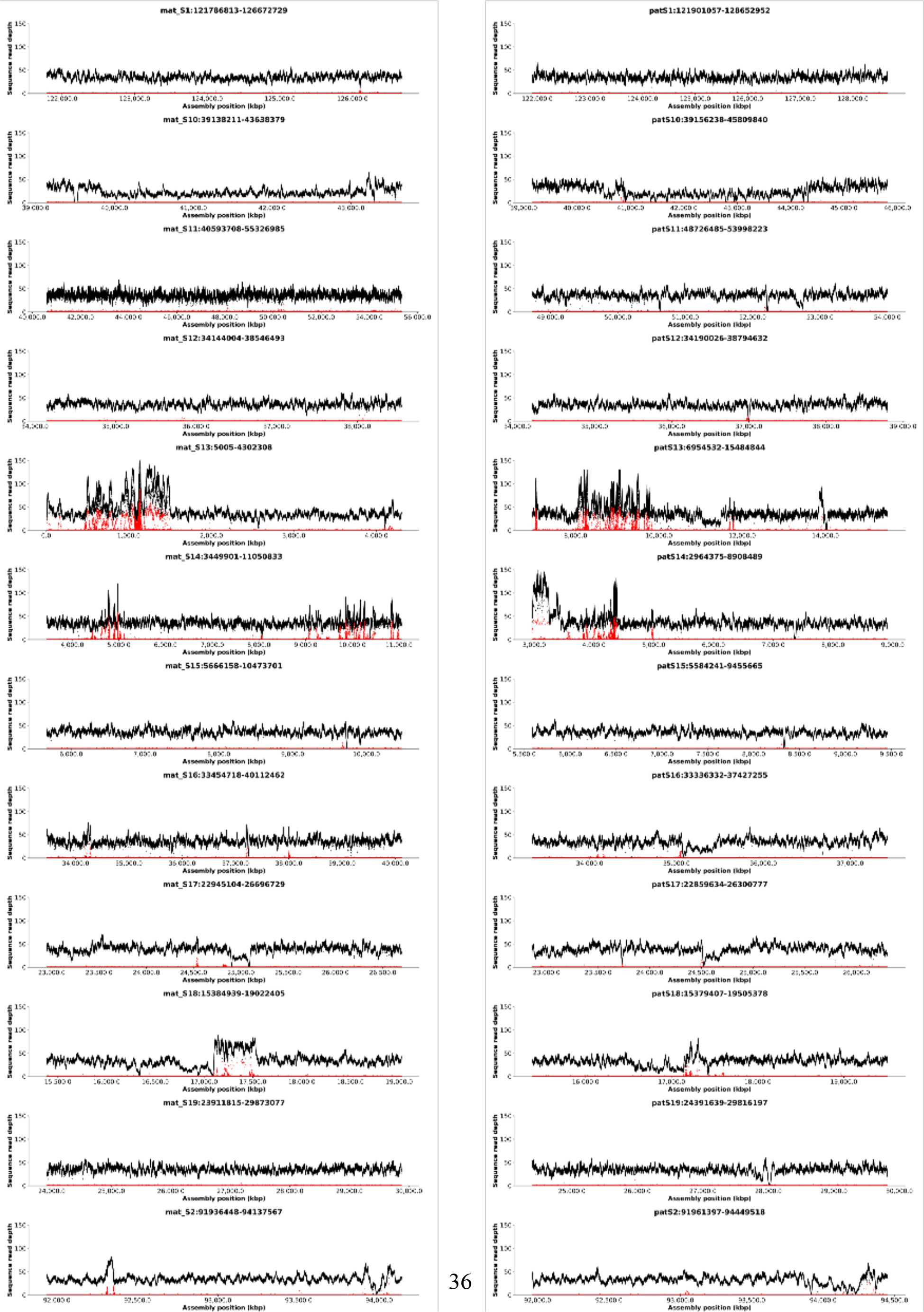
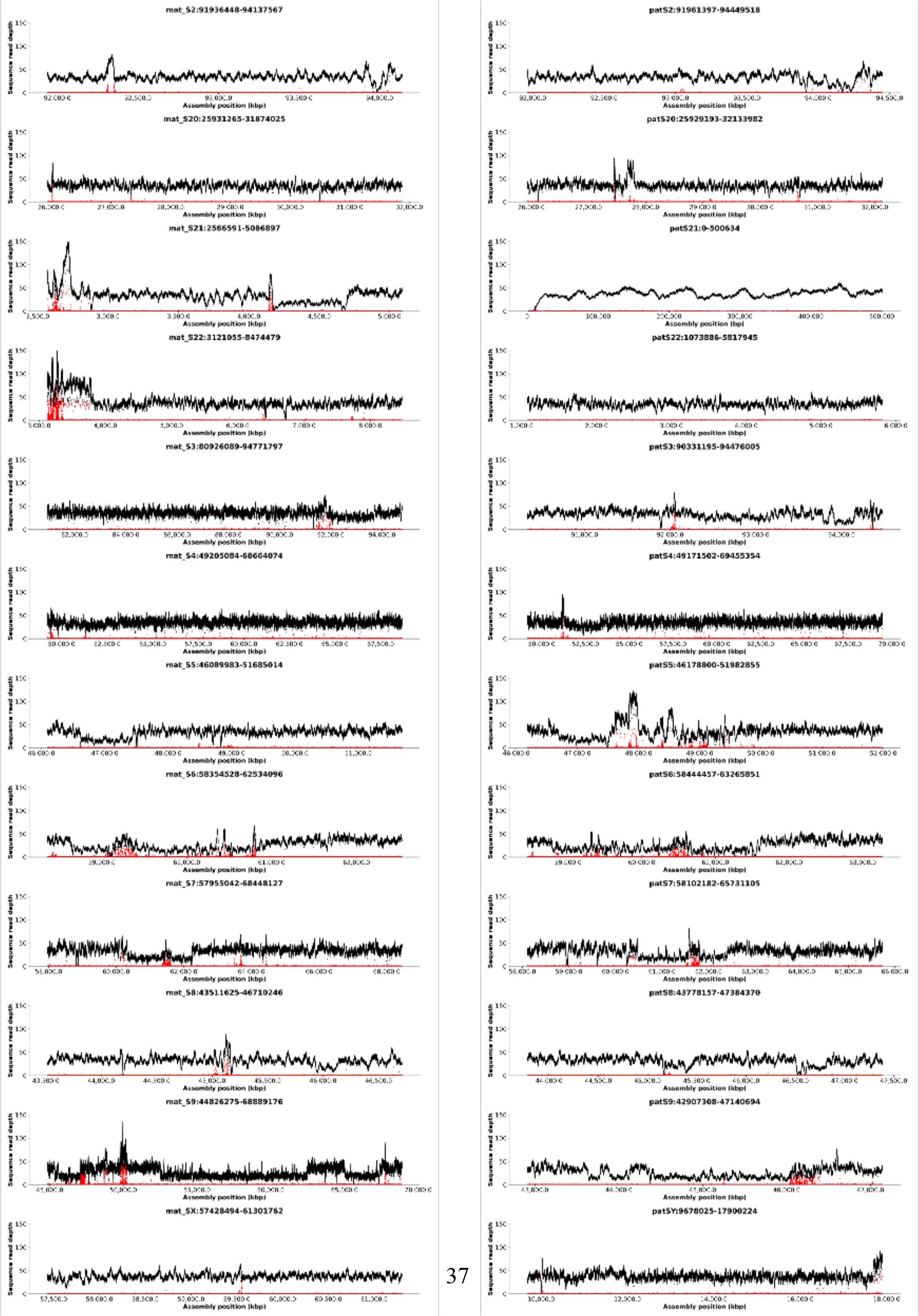
Assessing centromere collapses. Shown are HiFi coverage plots across the alpha satellite array regions of the centromeres in the HRPC-HG002 v1.0 maternal (left) and paternal (right) assemblies. A collapsed repetitive region will have coverage pile up that is two or more times higher than the mean coverage of the surrounding genomic region. These regions can also include alleles from the other haplotype (red), which could result in switch errors. Regions of decreased coverage or dropout in coverage could also have switch errors associated with them or are more difficult to sequence through.

**Supplementary Fig. 5.**
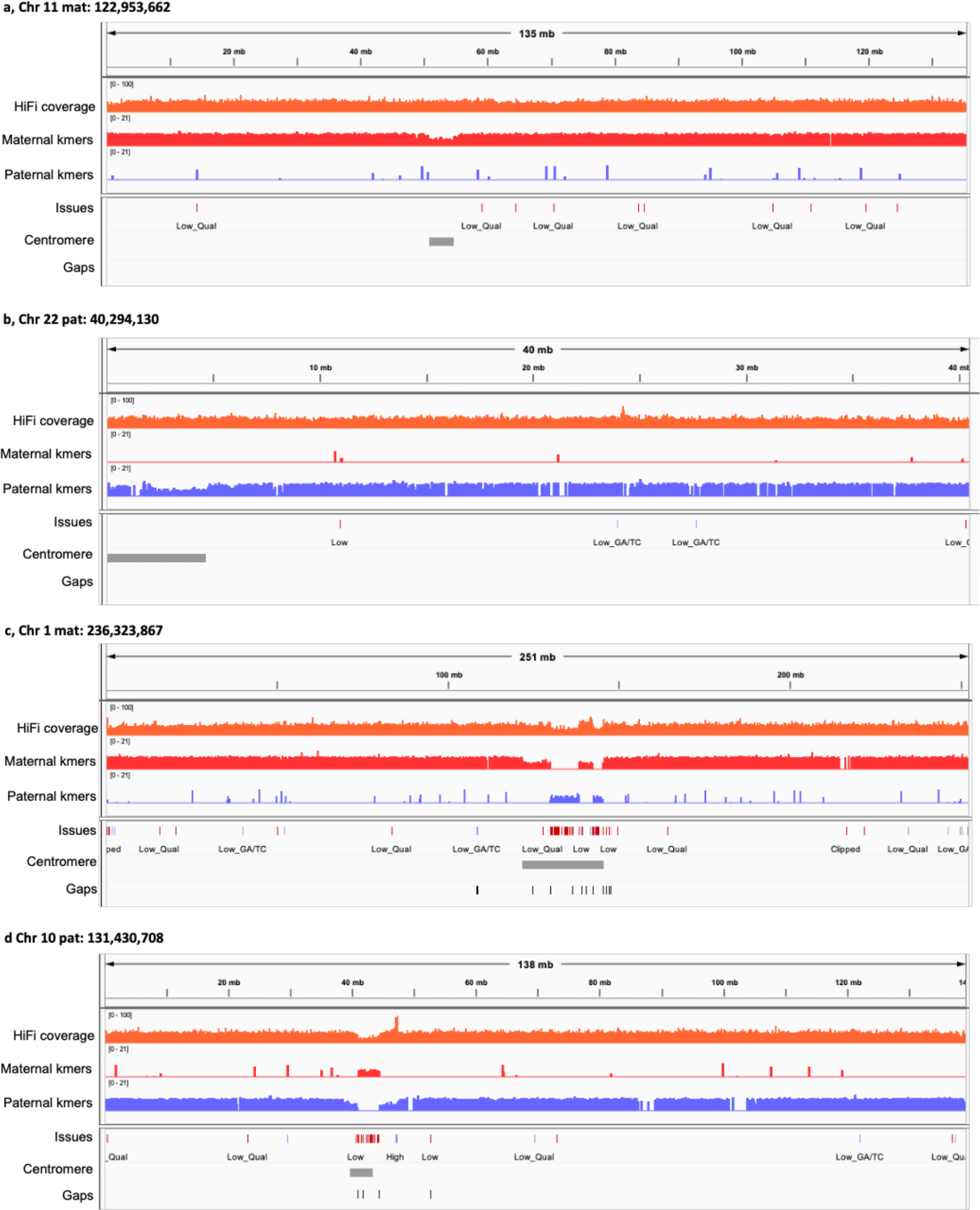
Assessing centromere switch errors. **a-b**, Examples of assembled HPRC-HG002 v1.0 centromere regions with no gaps or haplotype switch errors. **c-d**, Examples with gaps and haplotype switch errors. Top row, HiFi coverage across the chromosome segment; below that is maternal (red) and paternal (blue) *k-mer* profiles, where haplotype switches show up as swapping in k-mer coverage. Issues, indicate low quality sequence or other potential problems. Grey bar, centromere. Gaps, are black tick marks.

## Notes

https://github.com/human-pangenomics/HG002_Data_Freeze_v1.0

https://s3-us-west-2.amazonaws.com/human-pangenomics/index.html?prefix=

### Summary of Updates

Added two co-authors (Granat, Jaeger), corrected address of one co-author (Howe)

